# Bile acids target an exposed cavity in the glucocorticoid receptor modulating receptor self-assembly, chromatin binding and transcriptional activity

**DOI:** 10.1101/2025.05.13.653693

**Authors:** Alba Jiménez-Panizo, Thomas A. Johnson, Kaustubh Wagh, Andrea Alegre-Martí, Inés Montoya Novoa, Agustina L. Lafuente, Ulrich Eckhard, Luis Ángel Rodríguez-Lumbreras, Le Hoang, Martín Stortz, Montserrat Abella, Ido Goldstein, Annabel Valledor-Fernández, Lyuba Varticovski, Irwin Arias, Diego M. Presman, Diana A. Stavreva, Juan Fernández-Recio, Pablo Fuentes-Prior, Gordon L. Hager, Eva Estébanez-Perpiñá

## Abstract

The glucocorticoid receptor (GR) is an essential transcription factor that controls metabolism and homeostasis. Glucocorticoids (GCs) activate the GR upon occupying the internal ligand-binding pocket (LBP) of its ligand-binding domain (GR-LBD), which has been the focus of most previous structure-function studies. Synthetic GCs such as dexamethasone are widely used to treat inflammatory diseases, but their chronic use results in major side effects, whose molecular underpinnings remain unresolved. Here we present a thorough analysis of the topography of GR-LBD and its ability to bind small-molecule compounds, especially cholesterol derivatives. We show that one important class of steroids, bile acids, bind to previously unidentified and highly conserved, surface-exposed cavities on GR-LBD. We show that bile acids affect GR turnover and self-assembly in living cells, modulating receptor transcriptional activity. These findings reveal a previously unrecognized mechanism of GR regulation, with implications for the design of GCs with novel mechanisms of action.

**Teaser:** Bile acids modulate the activity of the glucocorticoid receptor upon binding to an exposed allosteric pocket thereby influencing transcriptional regulation and receptor self-assembly in living cells.

## INTRODUCTION

Nuclear receptors (NRs) tightly control a large spectrum of biological processes such as cell proliferation, metabolism, and homeostasis^1–3^; and are therefore one of the major pharmaceutical targets for many conditions including various cancer types, as well as inflammatory and metabolic diseases^4–9^. Because of their extraordinary pharmacological relevance, structure-function investigations of NRs have almost exclusively focused on their hormone- or ligand-binding pockets (LBPs), internal cavities within the ligand-binding domains (LBDs) mostly lined by aliphatic/aromatic residues^10–13^. However, evidence accumulated over the last two decades indicates the presence of surface-exposed pockets in several nuclear receptors that accommodate small molecules including endogenous hormones, metabolic compounds, and pharmacologically active molecules^14–18^.

Steroid receptors are a subfamily of NRs that bind with high affinity to a plethora of cholesterol-derived molecules collectively known as steroids^19^. The subclass includes the closely related receptors for glucocorticoids (GR; termed NR3C1 in the systematic nomenclature) and mineralocorticoids (MR; NR3C2), together with four members that control physiological processes of the reproductive system: the androgen receptor (AR; NR3C4), the progesterone receptor (PR; NR3C3), and estrogen receptors α (ERα; NR3A1) and β (ERβ; NR3A2)^1^.

The binding of pharmacologically relevant small-molecule compounds outside the LBP was first described for estrogen receptors^14,20^. The crystal structure of ERβ-LBD bound to 4-hydroxy-tamoxifen, the most abundant active tamoxifen metabolite, revealed the inhibitor occupying not only the LBP but also a surface groove that represents the major binding site of NR coregulators, also known as activating function 2 (AF-2)^20–24^. This feature may account for the mixed agonist/antagonist activity of type I antiestrogens such as tamoxifen and might explain its extraordinary success in the treatment of ER-positive breast cancer (recently reviewed in ref. ^25^). Nonsteroidal anti-inflammatory drugs and thyroid hormones that bind the AF-2 of AR have been identified as potential antiandrogens^16^. Finally, the AF-2 clefts of receptors from other subfamilies such as the thyroid hormone receptor α (TRα; NR1A1)^15^ and the vitamin D receptor (VDR; NR1I1)^17^ are also targeted by small molecules.

In addition to AF-2, we have previously identified a contiguous, solvent-exposed surface on AR-LBD that binds multiring compounds including thyroid hormones, called binding function 3 (BF-3)^16^. Of particular note, the AR BF-3 is a docking site for chaperones and co-chaperones^16,26–28^. Binding of a non-steroidal ligand to BF-3 inhibited AR by preventing its hormone-dependent dissociation from the Hsp90:FKBP52 complex^26^. This site is also conserved across NR subfamilies and suitable for drug intervention^16,29^. For instance, BF-3 regulates interactions with the linker between the DNA-binding domain and LBD of HNF4α (NR2A1)^30^. Finally, small-molecule compounds that bind outside the LBP have been reported in other receptor subfamilies, most notably for TRα^15,31^ and Nur77 (NR4A1)^32^, highlighting alternative modes of receptor modulation beyond the conventional ligand-binding paradigm. Despite substantial evidence indicating that well-defined surface cavities on the LBDs of nuclear receptors serve as functional binding sites with high druggability potential^18^, a thorough and systematic exploration of their structural topography remains limited. Moreover, co-crystal structures have remained an underutilized resource in drug discovery, with their valuable insights often overlooked in the search for new mechanisms of action and novel therapeutic targets.

Bile acids are a class of endogenous cholesterol-derived steroidal molecules that play essential physiological roles in nutrient absorption^33–35^, metabolism^36–39^, autophagy^40^ and longevity^41^. In addition to their well-characterized, non-specific activities as emulsifiers, bile acids are gaining increased attention as important signaling molecules in physiological pathways including hepatic lipid and energy metabolism, inflammation and immune homeostasis^42–44^, with important implications for disease^45–52^. These signaling activities of bile acids are mediated, at least partially, through their specific LBP-mediated interaction with farnesoid X receptor (FXR; NR1H4) together with liver X receptors α and β (LXRα; NR1H3 and LXRβ; NR1H2, respectively)^53–59^, as well as NR1I receptors, VDR^17,60,61^, pregnane X receptor (PXR; NR1I2)^62,63^, and constitutive androstane receptor (CAR; NR1I3)^64,65^. Accordingly, these members of the NR1H and NR1I subfamilies are collectively termed bile acid-activated receptors (BARs)^44,66^.

The gut microbiota plays a pivotal role in bile acid metabolism by converting primary into secondary bile acids. Primary bile acids, such as cholic acid (CA) and chenodeoxycholic acid (CDCA), are synthesized in the liver from cholesterol, conjugated with taurine or glycine, and secreted into bile, from where they are released into the intestine^67–69^. Once in the intestine, gut bacteria modify bile acids through enzymatic processes such as deconjugation and dihydroxylation^39,70,71^. These microbial transformations are crucial for maintaining the size and composition of the bile acid pool, which in turn influences hepatic and intestinal signaling pathways, including those mediated by FXR^72,73^. The glucocorticoid receptor has been connected to the regulation of inflammatory and immunomodulatory effects of bile acids such as ursodeoxycholic acid (UDCA)^74^. In addition, GR but not FXR modulates the therapeutic potential of the taurine-conjugated UDCA derivative, tauroursodeoxycholic acid (TauroUDCA), in preclinical models of the neurogenerative disorder, spinocerebellar ataxia type 3^75^.

Here we present a detailed analysis of the molecular topography of the ligand-binding domains of GR and other steroid receptors compared to that of canonical BARs. We identify common and unique cavities with ligand binding capabilities similar to those of the LBP and AF-2 pockets. GR, in particular, features a unique cavity predominantly lined by residues from helices H9/H10, which is important for receptor self-assembly and transcriptional activity *in vivo*. We explore the consequences of bile acid binding to this previously uncharacterized surface pocket on GR-LBD, demonstrating that it represents an actionable target for modulating both acute and long-term exposure to glucocorticoids. This binding event has significant implications for the glucocorticoid receptor function and offers potential avenues for therapeutic intervention.

## RESULTS

### The hormone-binding domain of the glucocorticoid receptor features solvent-exposed cavities with high ligand-binding potential and significant druggability

To explore the topography of the LBD modules from GR and other steroid receptors, we conducted a thorough bioinformatics analysis using state-of-the-art programs for pocket detection and druggability potential^76–78^. For comparison, we also analyzed the LBDs of two BARs, FXR and VDR, for which structural evidence of bile acid binding has been reported^17,58,60^. The results of these analyses are presented in Fig. 1A, Fig. S1A and Tables S1 to S3. As expected, the well-characterized LBP and AF-2 pockets of all receptors are identified with high ligand-binding abilities and medium to strong druggability. However, we observed large differences regarding other surface cavities. Most notably, all programs consistently identify a major cavity in the GR-LBD surface lined by residues from helices H9/H10 and the connecting loop, L9-10, with a higher ligand-binding ability than the LBP and a medium druggability score (Table S1). The previously characterized BF-3 site is also identified in this systematic unbiased analysis, but with weak druggability (Fig. 1A).

**Fig. 1.**
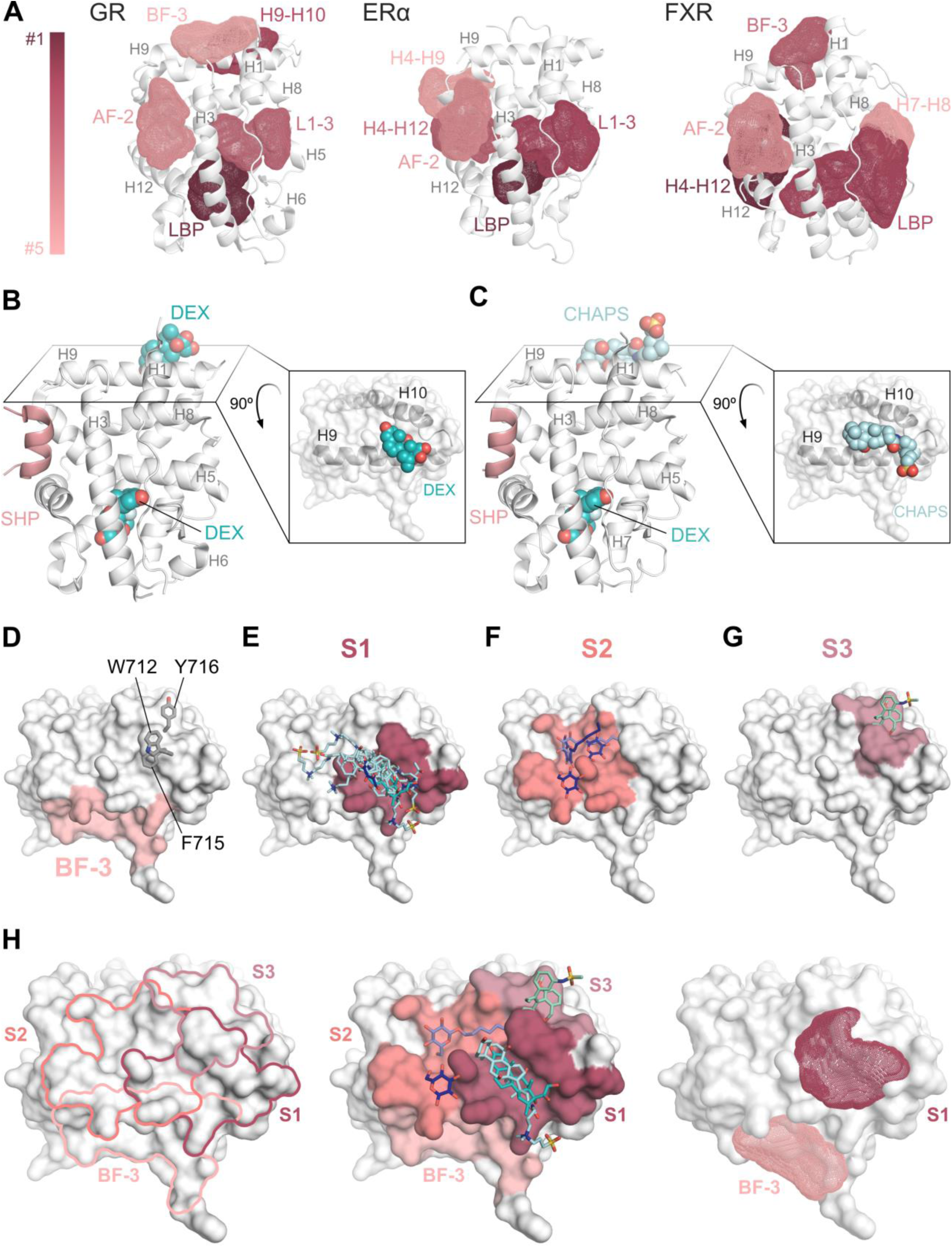
The ligand binding domain of GR and other nuclear receptors display several well-defined solvent-exposed cavities. **A**, Representation of the top five cavities in the LBD of two steroid receptors, GR and ERα, as compared to FXR. Major helices and loops are labeled. Pockets are colored based on their druggability scores. Note the unique cavity network of GR-LBD. **B, C**, Small-molecule compounds such as Dex or CHAPS bind to a GR-LBD cavity formed by helices H9/H10 and the connecting loop, L9-10. GR-LBD is depicted as a white cartoon and bound small molecules are shown as color-coded sticks (PDB codes 7YXP and 7YXO, respectively). **D**, Surface representation of the GR domain viewed from the top in the standard orientation. The BF-3 site is highlighted in salmon. Aromatic residues at the N-terminus of H10 are shown as sticks and labeled. **E-G**, Superimposition of all compounds that bind at the major surface-exposed cavity of GR-LBD in deposited crystal structures. Depending on the contact residues, three different subsites can be defined. (**E**) The most frequently targeted subsite S1 includes residues from the more C-terminal end of H9, the L9-10 loop, and Trp712 / Phe715 in H10. Bound BOG, Dex and CHAPS molecules (from PDB codes 1M2Z, 7YXP, 4UDD and 7YXO, respectively) are shown as colored sticks, with cyan carbon atoms. (**F**) Subsite S2 features residues from the N-terminal half of H9, Phe715 / Thr719 in H10, and extends towards the C-terminal F domain (e.g., Phe774). BOG and JZR molecules (PDB codes 1M2Z and 3K22, respectively) are indicated as color-coded sticks with dark blue C atoms. (**G**) Subsite S3 binds the LSJ molecule (PDB 4LSJ) - represented as color-coded sticks with pale green C atoms - and includes the triplet of aromatic residues at the N-terminus of H10 together with His775. **H**, Relationships between BF-3 and S subsites at the top interface of GR-LBD. The boundaries of the subsites are shown in the left panel, colored as in figures 1D-G. The four surfaces are shaded in the central panel, with selected compounds shown as color-coded sticks. Note that subsite S1 matches the computationally identified cavity that is only second to the LBP in terms of druggability (left panel). (See Fig. S1 for the surface topography of other NRs and Table S1 for more details).

### A surface-exposed cavity lined by helices H9 / H10 and the connecting loop is a common binding site for small molecules

Given the high ligand affinity and significant druggability predicted for some solvent-exposed cavities in GR-LBD, we explored the crystallographic evidence that supports their capacity to bind small molecules. In our recently reported structures of GR-LBD, we unexpectedly observed extra electron density at the ‘top’ homodimerization interface of the domain, which could be interpreted as either the commonly used bile acid-derived detergent, CHAPS, or as a Dex molecule (PDB entries 7YXO and 7YXP^13^) (Fig. 1, B and C). The observation of these cholesterol-derived compounds bound to a BF-3–proximal surface prompted us to systematically review deposited crystal structures for small molecules bound to the GR-LBD surface. The results of this analysis are summarized in Fig. S1, B to D, and reveal three different subsites that overlap with the ‘top’ interface of GR (labeled S1 to S3 in Fig. 1, E to H)^13^. Notably, the conformation of the Trp712 indole moiety varies to accommodate these small molecules in the different crystal structures (Fig. S1, B and C). Subsite S1, which corresponds to the most prominent cavity identified in our bioinformatics analyses (Tables S1-S3), is a major binding site for molecules featuring from one to three rings. This cavity exhibits notable flexibility and adaptability, as evidenced by variations in the solvent-accessible surface area across the various crystal structures (Fig. S2A). Moreover, the electrostatic potential of the GR-LBD surface reveals that this region is predominantly neutral, providing an optimal environment for ligand docking through hydrophobic interactions (Fig. S2B). Because binding to this solvent-exposed groove could regulate GR biological activity in living cells (see below), we designated the subsite S1 as ‘sensor function-3 (SF-3)’, with S2 and S3 as lateral extensions of this cavity. SF-3 comprises residues from helices H9/H10 and the connecting loop, most notably a triplet of conserved aromatic residues at the N-terminus of H10 (Trp712, Phe715 and Tyr716; Fig. 1D). SF-3 is thus contiguous to but topologically distinct from the BF-3 pocket^15,29^. Another noticeable surface cavity formed by neighbouring residues of the L1-3 loop (e.g., Trp557), L6-7/H7 (e.g., Met639, Cys643), and helix H11 (most notably Tyr735) was also identified as binding site for small molecules in several crystal structures. This pocket has been postulated as a hormone entrance channel to the LBP in several NRs^79^, and therefore will be termed entry/exit (E) site in the following (Fig. S1D).

Several observations from previous crystal structures solved at higher resolution are particularly noteworthy. (1) In addition to CHAPS, molecules of other non-ionic detergents, octyl β-D-glucoside (BOG) and hexyl β-D-glucoside (HOG), occupy SF-3 in entries 1M2Z^10^ and 3K22^80^, respectively (Fig. S1B). (2) Some GR-LBD modules feature up to three bound CHAPS molecules. One of these detergents, which overlaps well with the multiring ligands found in our crystals, docks on helix H9 with its head trapped between the aromatic side chains of Trp712 and Phe715 (4UDD^81^) (Fig. S1B). A second binding site for CHAPS is observed in this structure and in entries 4UDC, 5G3J^82^, and 5NFP^83^, which overlaps with the E site (Fig. S1C and D). Finally, and particularly relevant for its pharmacological implications, (3) two molecules of the nonsteroidal GC, LSJ, occupy both the LBP and SF-3 in entry 4LSJ (Fig. S1B); the latter making strong contacts with the Trp712 / Phe715 / Tyr716 triplet^84^. Interestingly, temperature factors for both GC molecules are similar (average B factors 22.7 vs. 27.8 Å^2^), highlighting the strong binding of the compound to SF-3. Overall, while ligand binding relies mostly on Van der Waals (VdW) contacts, several polar/charged residues in SF-3 can engage in electrostatic interactions (e.g., Ser708 and Asn711), providing specificity for ligand recognition.

The SF-3 site participates in important homo- and heterotypic interactions. Most notably, SF-3 residues are involved in several of the 20 possible arrangements of GR-LBD homodimers in our recently presented catalog^13^. This is in addition to several symmetric, parallel homodimers in which the H10 aromatic triplets of two monomers contact each other^85^, thus effectively juxtaposing their SF-3 pockets. The gathering of the six exposed aromatic side chains (Trp712/712’, Phe715/715’, and Tyr716/716’) generates large cavities with potential higher ligand binding capacity. In fact, head and tail of the best-defined BOG molecule in 1M2Z occupy the SF-3 pockets of neighbouring monomers. These observations suggest that occupancy of SF-3 by small-molecule compounds could modulate self-assembly of the full-length receptor. Furthermore, the recent cryo-EM structure of the human GR-Hsp90-p23 complex reveals that p23 directly interacts with this hydrophobic patch, stabilizing native GR and enhancing hormone binding (Fig. S1E)^28^.

### Analog screening identifies bile acids as preferred SF-3 ligands

To further explore the binding potential of the solvent-exposed SF-3 pocket, we performed an analog search across the 8 million compounds deposited in the ZINC and DrugBank databases, focusing on structural and chemical similarities to cholesterol-derived molecules. Using both fingerprint and substructure-based approaches, we identified 36 Dex and CHAPS analogs as potential GR-LBD ligands (Table S4). The high prevalence of bile acids among the identified molecules suggests that the SF-3 cavity is a preferred binding site for these cholesterol derivatives. In addition, analog searches guided by the bound BOG molecules identified several polyphenols with known anti-inflammatory activity as potential SF-3 interacting molecules (Fig. S2C and Table S4).

Analogs for which structures were available were docked into the SF-3 cavity of human GR-LBD (PDB 4UDD) using different constraints. Lithocholic acid (LCA) and other bile acids fitted perfectly into SF-3 without clashes with the protein (Fig. 2, A to C). Notably, three of the top analogs selected for the presence of a hydrogen bond with the Ser708, as seen in the crystal structure, formed additional salt bridges between carboxylate or sulfonate moieties of the bound bile acid and the Arg690 guanidinium group, underscoring their strong electrostatic complementarity to SF-3. These observations suggest that conjugated bile acids with longer, acidic tails might bind more tightly and with higher specificity to this cavity. Docking experiments with AlphaFold3 (Fig. S2D) and AutoDock (Fig. S2E) suggested that LCA could dock inside the LBP of human GR-LBD, with only minor clashes with residues such as Met604 and Phe623. However, most AutoDock runs locate the bile acid outside the LBP. Furthermore, closer inspection of these docking poses reveals that LBP-bound LCA molecules would not be able to engage in H-bonds with the Gln570 carboxamide group as in crystal structures of Dex/Cort-bound GR-LBD (Fig. S2F). Binding of conjugated bile acids such as taurocholic acid (TauroCA) inside the LBP would be even less favorable, as it would require substantial rearrangements of the LBP, and no such docking solutions were found (Fig. S2, G to I). Of note, AlphaFold3 predictions of TauroCA binding to human GR-LBD in the presence of Cort show the E and SF-3 cavities as preferred binding sites for the bile acid. Predicted binding energies of these interactions calculated by pyDock scoring suggest a more favorable ligand binding to SF-3 compared to the E site (-12.0 vs. -9.9 kcal/mol, Fig. S2J).

**Fig. 2.**
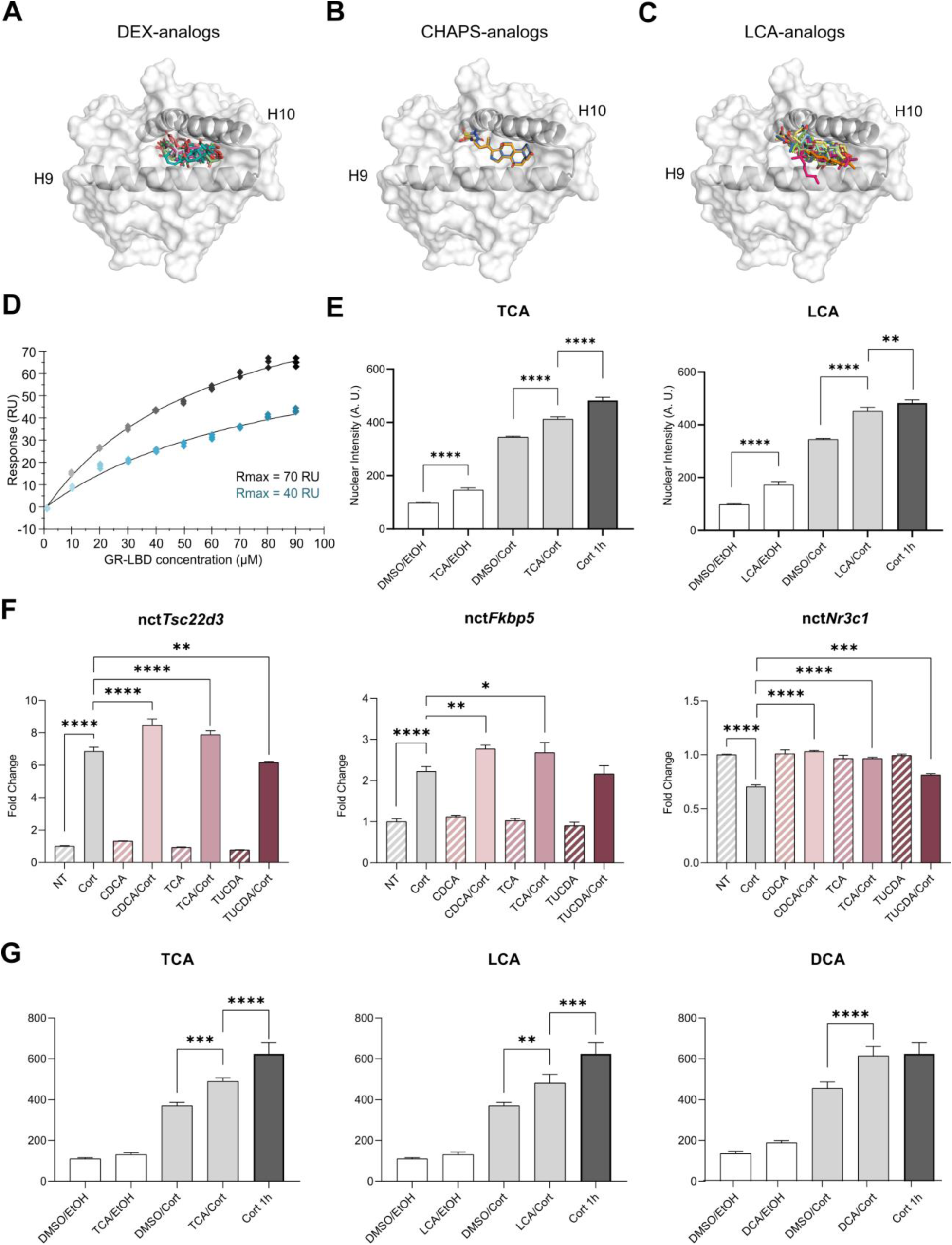
Bile acid binding to GR-LBD reduces self-association in solution, reverts the Cort-mediated negative feedback loop in cells and protects GR from proteasomal degradation. **A-C**, Several multiring compounds, most notably bile acids, were identified as SF-3 binders in high-throughput docking experiments conducted using either Dex (**A**), CHAPS (**B**) or LCA (**C**) as templates. The top compounds identified in these screens are represented as colored sticks. **D**, LCA binding to GR-LBD interferes with protein self-association in solution. SPR experiments were performed by running increasing concentrations of Dex-bound GR-LBD (0.2, 0.4, 0.8, 1.6, 3.12, 6.25, 12.5 and 25 µM) in the presence or absence of LCA over chip-immobilized GR-LBD. The fitted curves correspond to duplicate experiments conducted in the absence (black) or presence (blue) of 40 µM LCA. Protein-protein self-association was analyzed according to a 1:1 model. The results of assays conducted in triplicate are shown along with the respective maximum response. **E**, Nuclear intensity levels of endogenous GR after bile acid incubation for 24 h in 3134 cells assessed by immunofluorescence staining (Ab: G-5). Cotreatment with 100 nM Cort and 100 µM TCA or LCA leads to a significant increase in nuclear GR protein compared to vehicle. Statistical analysis was performed using one-way ANOVA followed by Tukey’s post-hoc test to compare group means. Significant differences at p values < 0.05 are denoted by asterisks (* p < 0.05, ** p < 0.01, *** p < 0.001 and **** p < 0.0001). **F**, Impact of 24 h treatment with 100 µM of the indicated bile acids on GR transcription, assessed by RT-qPCR. Mean relative expression levels are normalized to a housekeeping gene (*18-S*), with error bars indicating standard deviations from three biological replicates. Statistical analysis was performed using one-way ANOVA followed by Tukey’s post-hoc test to compare group means. Significant differences at p values < 0.05 are denoted by asterisks (* p < 0.05, ** p < 0.01, *** p < 0.001 and **** p < 0.0001). **G**, Nuclear intensity levels of Halo-TMR ligand-stained endogenous Halo-GR in 3134 cells was measured after cotreatment of Cort and bile acid for 24 h and compared to Cort/DMSO (24 h). As an additional control, untreated Halo-TMR-stained cells were cultured for 24 h in serum-free medium and activated with Cort for 1 h to account for the loss of ligand-stained Halo-GR due to the culturing in a serum-free medium (Cort 1 h). Statistical analysis was performed as described above (E).

### Bile acids and non-ionic detergents bind to immobilized GR-LBD and interfere with receptor self-association in solution

To explore the possible functional relevance of small-molecule binding to GR-LBD and its implications for receptor multimerization, we performed surface-plasmon resonance (SPR) assays to verify whether cholesterol-related compounds and other non-ionic detergents might directly influence homotypic interactions in solution (see Fig. S3A for the chemical structures of CHAPS and tested bile acids). First, we verified that detergents CHAPS and BOG bind to immobilized, Dex-bound GR-LBD with micromolar affinity (Fig. S3, B and D). Next, we assessed whether these compounds could modulate GR-LBD protein-protein interactions in solution. To this end, we conducted further SPR experiments in which the binding sites in the immobilized GR-LBD were first saturated with detergent (1 mM CHAPS or BOG). Then, increasing concentrations of GR-LBD were run over the Biacore chip, also in the presence of buffers supplemented with the same saturating detergent concentrations. Indeed, saturation of CHAPS binding sites led to a two- to three-fold decrease in the strength of protein-protein interactions, depending on the model used to fit SPR data, when compared to our previous results for WT GR-LBD^13^ (Fig. S3C). By contrast, no significant effects on the 1:1 interaction were detected in the presence of BOG (Fig. S3E). These results demonstrate that cholesterol-derived detergents such as CHAPS can bind GR-LBD, likely to SF-3, and interfere with GR-LBD self-association in solution. Importantly, two physiologically relevant bile acids, LCA and TauroCA, bound to GR-LBD with higher affinity than CHAPS (k_D_ = 4.9 and 6.0 µM, respectively; Fig. S3, F and G). Furthermore, incubation of saturating concentrations of LCA interfered with GR-LBD self-association, reducing the maximum response by over 40% (Fig. 2D).

### Long-term treatment with bile acids protects GR from hormone-induced repression and proteasomal degradation

Given the structural and computational evidence indicating that bile acids are preferred SF-3 ligands and have an impact on GR-LBD self-association in solution, we decided to explore their potential impact on the turnover of the full-length receptor. The SF-3 pocket is recognized by the Hsp90-p23 chaperone complex, which plays an essential role in maintaining the folded state of GR, preventing its premature degradation^28^. E3 ligases like CHIP recognize and ubiquitinate free GR, which is followed by its proteasomal degradation^86,87^. Serum bile acid concentrations in humans can vary significantly based on factors such as age, gender, and health conditions. In healthy individuals, total serum levels are typically below 5 µM. However, in certain pathological conditions, bile acid levels can rise substantially, reaching concentrations over 100 µM^88–90^. Intracellular levels of bile acids are more challenging to measure and can vary significantly between different cell types and pathophysiological conditions^91^.

To assess whether bile acids can interfere with this process, we treated 3134 mouse mammary adenocarcinoma cells with 100 nM Cort in the presence or absence of 100 µM non-conjugated (deoxycholic acid; DCA, LCA, CDCA) or taurine-conjugated bile acids, (TauroCA, TauroUDCA) for 24 h and measured the expression levels of endogenous GR by immunofluorescence staining. GR protein levels were higher both in the nucleus and the cytoplasm compared to untreated controls (Fig. 2E and S4A, B). We also detected an increase in the expression levels of nascent *Fkbp5* and *TTscc22d3* genes after bile acid treatment for 24 h (Fig. 2F), indicating that the higher GR concentrations affect its transcriptional outcome. To discern whether the effects of long-term treatment with bile acids are due to protection from protein degradation or because of an increase in GR transcription, we performed qPCR analysis of nascent *Nr3c1* under the same conditions. The results show a decrease in GR expression after 24 h of Cort induction (Fig. 2F), which is in line with previous observations of an inhibitory feedback loop triggered by prolonged hormone activation^92–94^. The observed Cort-repression was reverted when co-treating with bile acids for 24 h, suggesting that these cholesterol-derivatives counteract GC-triggered inhibitory feedback mechanisms.

To assess whether bile acids could independently affect GR protein stability, we performed additional experiments with cells expressing Halo-tagged endogenous GR essentially as described above. Labeling with the specific fluorescent HaloTag ligand, TMR, enabled us to selectively track pre-existing receptor molecules, thus avoiding interference from newly synthesized protein. Quantification revealed a significant increase in TMR signal in cells treated with LCA, DCA or TauroCA compared to untreated controls (Fig. 2G), indicating that bile acid treatment protects GR from degradation over the 24-hour period. Taken together, our results suggest that long-term incubation with bile acids not only interferes with the hormone-induced negative feedback that typically represses GR gene expression but also stabilizes GR protein by preventing its proteasomal degradation, demonstrating dual modulation at both transcriptional and post-translational levels.

### Short-term treatment with bile acids downregulates GR transcriptional activity

Long-term treatment with both conjugated and non-conjugated bile acids potentially inhibits Cort-induced repression of *Nr3c1* expression (Fig. 2E-F and Fig. S4). Therefore, we decided to further explore their impact on the short-term transcriptional activity of GR. We first tested whether these cholesterol-like compounds could affect early human GR transcriptional activity after GC induction. For that, we treated human leukemic monocyte THP1 cells with 10 nM Dex, alone or in combination with 20 μM CHAPS or LCA. As expected, Dex potently induced expression of *Tsc22d3*, when compared to non-stimulated cells. However, this effect was suppressed in the presence of the cholesterol derivatives (Fig. S4C). Subsequent experiments were carried out with 3134 mouse mammary adenocarcinoma cells, as they were also used for assessing GR dynamics and function in living cells (see below).

We assessed the impact of three secondary bile acids on early GR function: LCA, the closely related UDCA, and TauroCA, with a longer tail and a terminal sulfonate group due to its conjugation with taurine (see Fig. S3A for the chemical structures of the tested compounds). Of note, concentrations of 100-200 µM of LCA and UDCA have previously been shown to possess anti-inflammatory activity in colonic epithelial cells and to confer protection from colonic inflammation *in vivo*^95^. We treated GR knock-out cells derived from the 3134-mouse mammary adenocarcinoma cell line (3134-GR^KO^) expressing GFP-tagged mouse GR with different concentrations of these bile acids for 2 h. Then, we analyzed the expression levels of several classical GR target genes (*Per1*, *Tsc22d3*, *Sgk1* and *Arld4*) by qPCR. To prevent excessive GR activation, we treated these GFP-GR expressing cells with a low, more physiologically relevant concentration of Cort (100 nM). Pre- and co-treatment with different concentrations of all three bile acids led to a significantly reduced expression of *Per1* and *Sgk1* genes, indicating an antagonistic effect on GR transcriptional activity under these conditions (Fig. 3A). This modulatory effect was not observed when higher Cort concentrations (400-600 nM) were used, or when cells were stimulated with 100 nM of the more potent, synthetic GC, Dex (Fig. S5).

**Fig. 3.**
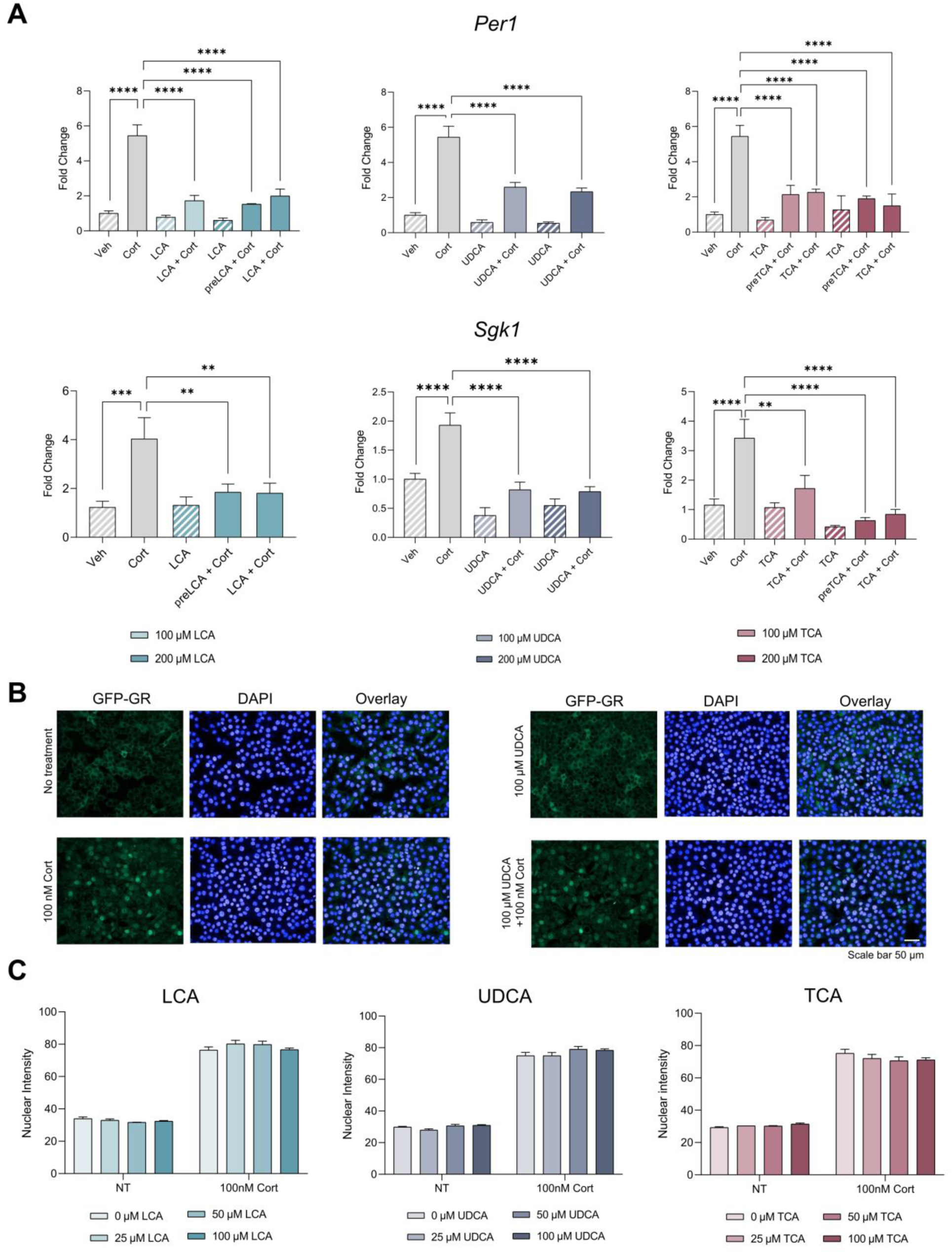
Bile acids downregulate GR transcriptional activity without affecting its nuclear trafficking. **A**, Effect of 2 h treatment with bile acids on the expression of two major GR-responsive genes, *Per1* (top) and *Sgk1* (bottom), as assessed by RT-qPCR. Mean relative expression levels are normalized to a housekeeping gene (*β-actin* or *18-S*), with error bars indicating standard deviations from three biological replicates. Statistical analysis was performed using one-way ANOVA followed by Tukey’s post-hoc test to compare group means. Significant differences at p values < 0.05 are denoted by asterisks (*p < 0.05, ** p < 0.01, *** p < 0.001 and **** p < 0.0001). **B**, Representative images of GFP-GR nuclear translocation in 3134 cells in response to 100 nM Cort, 100 μM UDCA, or both. Nuclei were stained with DAPI. Scale bar, 5 μm. **C**, Quantitative analysis of GFP-GR nuclear translocation upon incubation with either non-conjugated (LCA, UDCA) or conjugated bile acids (TauroCA, TCA). Translocation rate was calculated as the ratio of nuclear versus cytoplasmic intensity and each value was normalized to the control. Error bars represent the mean value ± standard error of the mean, n = 6.

### Bile acids do not influence GR trafficking to the nucleus

The negative impact of bile acids on GR function in living cells could result from their interference with GR translocation to the nucleus. To explore this possibility, we assessed the translocation efficiency of cells treated with LCA, UDCA or TauroCA for 1 h, as compared to unstimulated cells (Fig. 3, B and C). Bile acid treatment alone did not promote GR translocation to the nucleus, indicating that, unlike steroid hormones, these cholesterol derivatives do not occupy the ligand-binding pocket of the receptor in a productive conformation. These results further validate the docking experiments and current crystallographic evidence described above and are in line with bile acids binding to SF-3 in living cells. Furthermore, GR translocation induced by Cort was not affected by co-treatment with any of the tested bile acids (Fig. 3C). Thus, short-time treatment with bile acids interferes with GR activity specifically in the nucleus.

### Bile acids interfere with GR function at the chromatin level

To verify that bile acids affect GR once the receptor is translocated into the nucleus, we first compared the expression levels of *Per1* and *Sgk1* at various time points ranging from 0.5 to 3 hours after a 30 min activation with Cort alone or in combination with 100 µM TauroCA (see Figure 4A for a schematic representation of the experimental protocol). A significant reduction in expression levels was observed at the peak of induction for both genes in cells co-treated with TauroCA (*Per1* after 0.5 and *Sgk1* after 3 h), indicating that the bile acid rapidly downregulate GR activity and sustain this inhibitory effect over several hours (Fig. 4B). Then, we conducted the same experiment but adding TauroCA 30 min after Cort activation, thus ensuring a complete translocation of GR into the nucleus before bile acid treatment. The results of this time-course assay showed a similar reduction of *Per1* and *Sgk1* expression levels compared to previous conditions (Fig. 4C), corroborating that the bile acid primarily affects GR function after its localization to the nucleus.

**Fig. 4.**
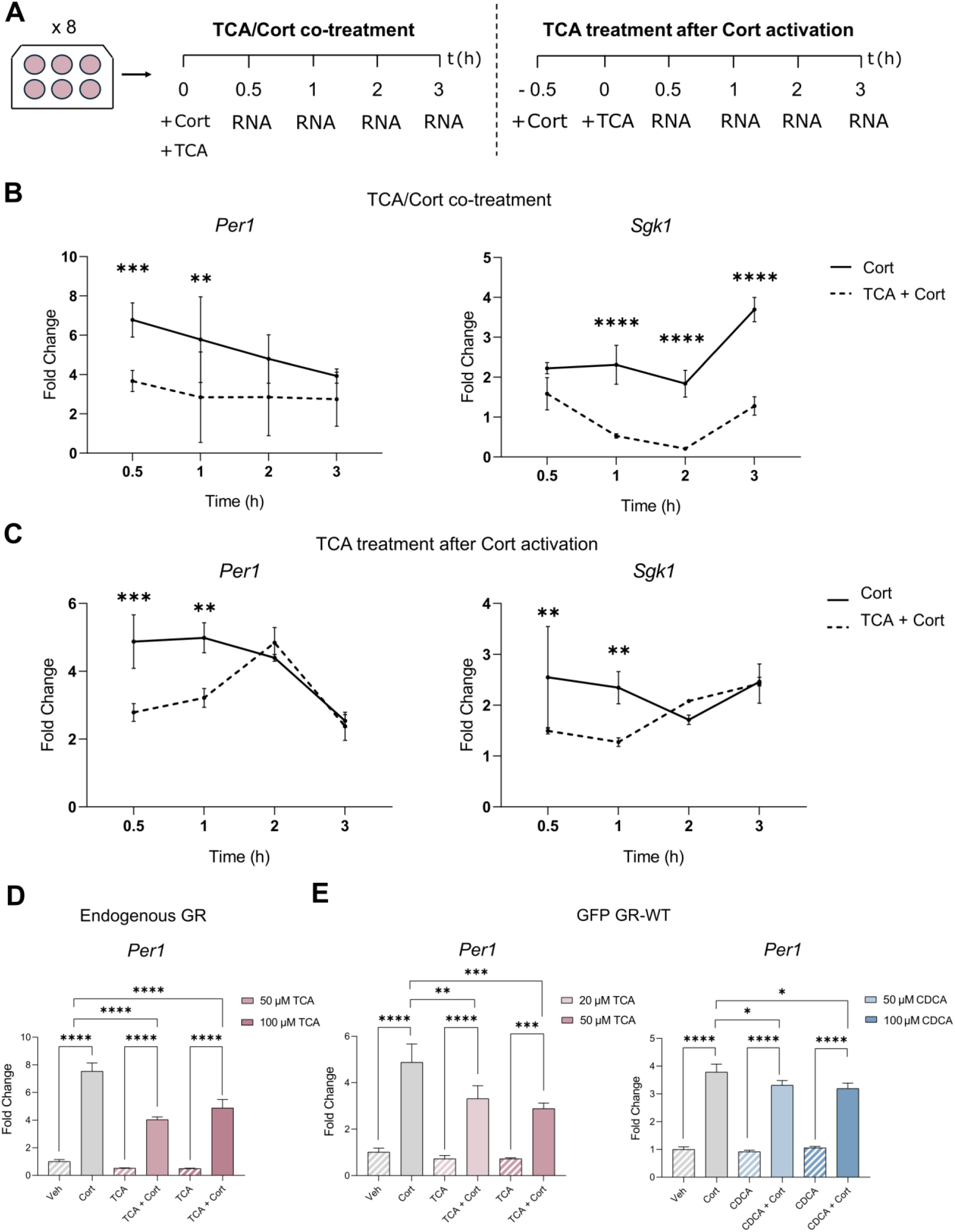
Bile acids specifically downregulate the nuclear activity of GR. **A, B**, Time course of *Per1* and *SGK* gene expression (from 0.5 to 3 hours) as assessed by RT-qPCR. Cells were either cotreated with 100 nM Cort and 100 μM TauroCA (TCA) (**B**) or treated first with the hormone for 30 min before bile acid addition (**C**). Note the faster induction of *Per1*, which shows a maximum response 30 min after induction in both scenarios. Note also that the bile acid effect is significant at the time points of maximum gene expression. **D**, Downregulation of *Per1* gene expression after 30-minute co-treatment of 3134 cells, expressing endogenous GR, with 100 nM Cort and either 50 or 100 μM TauroCA. **E**, *Per1* expression is downregulated in cells expressing GFP-tagged GR by co-incubation with 20 or 50 µM TauroCA and 50 or 100 µM CDCA. Mean relative expression levels for all the experiments shown in panels A-D were normalized to a housekeeping gene (*β-actin* or *18-S*), with error bars indicating standard deviations from three biological replicates. Statistical analysis was performed using one-way ANOVA followed by Tukey’s post-hoc test to compare group means. Significant differences (p < 0.05) are denoted by asterisks (*p < 0.05, ** p < 0.01, *** p < 0.001 and **** p < 0.0001).

Additional experiments using 3134 cells expressing endogenous GR demonstrated that the downregulatory activity of the bile acids was not conditioned by the GFP tag (Fig. 4D). Additionally, we assessed whether the observed effects were maintained at lower bile acid concentrations. Indeed, concentrations as low as 50 µM of the non-conjugated bile acid, CDCA, or 20 µM of TauroCA were able to significantly reduce *Per1* expression (Fig. 4E), demonstrating potent bile acid-mediated downregulation of GR transcriptional activity.

To further analyze the impact of bile acids on GR upon nuclear translocation, we conducted single-molecule tracking (SMT) experiments^96^ to follow the intranuclear dynamics of GR within living cells in the presence or absence of TauroCA. SMT has emerged as a cutting-edge technology for directly visualizing and tracking individual proteins in living cells, particularly transcription factors^97^. Recent studies emphasize the importance of considering both specific and non-specific protein-chromatin interactions when interpreting transcription factor movement and further transcriptional activity^97^. We performed SMT on 3134 cells expressing HaloTag-mGR or Halo-H2B chimeras (with H2B serving as a probe for photobleaching) to determine the spatial mobility of these proteins in the nucleus after 1 or 3 h treatment with 100 µM TauroCA. Since LCA can modulate GR oligomerization in solution (Fig. 2D), we performed SMT experiments in cells expressing HaloTag-mGR P481R (also known as GR^tetra^, a constitutively tetrameric GR mutant)^98,99^. As shown previously, GR-WT and GRtetra exhibit power-law distributed dwell times^100^. TauroCA significantly reduces the dwell time of WT GR after 1 hour of treatment (Fig. 5C), pointing to either a reduction in receptor stability when binding to chromatin or an increased dissociation rate. We observed that, in the absence of bile acid, the dwell times of WT GR and GR^tetra^ are comparable. However, no effect was observed for the tetrameric mutant after 1 hour of TauroCA treatment (Fig. 5, C to E). These results suggest that bile acids cannot modulate GR structure and function when receptor multimerization is enhanced (e.g., by modifying the inter-monomer interfaces). We also examined the spatial mobility of GR in the presence and absence of TauroCA and found GR to exhibit two distinct low-mobility states as described previously for GR and other chromatin associated proteins^101–104^. However, the higher mobility state was slightly reduced, without affecting the total bound fraction or the fraction of the lowest mobility state, which is preferentially occupied by activated GR (Fig. S6, A and B).

**Fig. 5.**
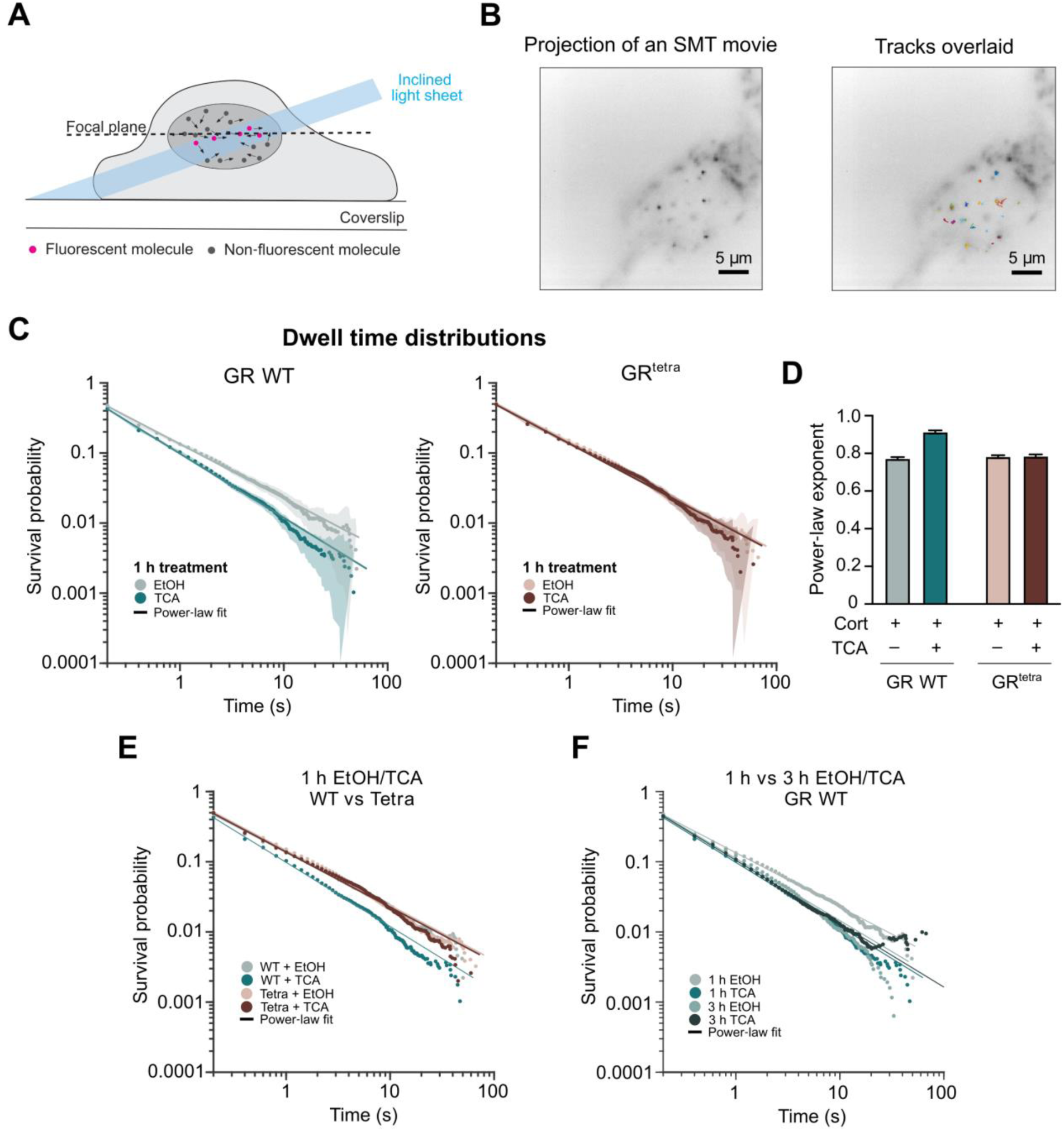
Survival distributions of GR dwell times upon bile acid treatment. **A**, Schematic representation of the SMT setup using highly inclined and laminated optical (HILO) microscopy. A thin section of the sample is illuminated, reducing background fluorescence and enhancing signal-to-noise for single-molecule detection. Fluorescent molecules within the focal plane (magenta dots) are selectively excited and tracked. **B**, Time projection of a representative Halo-GR WT SMT movie (left panel) and tracks generated by the observed GR molecules over time (right panel). **C**, SMT of Halo-WT GR and Halo-GR^tetra^ activated with 100 nM Cort (Ncells WT = 36, Ntracks WT = 9078, Ncells Tetra = 38, Ntracks Tetra = 18363), in the presence or absence of 100 µM TauroCA (Ncells WT = 42, Ntracks WT = 13916, Ncells Tetra = 36, Ntracks Tetra = 15482). Images were collected every 200 ms with an exposure time of 10 ms. The survival distributions are shown as solid lines, fitted to a power-law function. **D**, Power-law exponent of Halo-WT GR and Halo-GR^tetra^ under different treatments with Cort and/or TauroCA. Note that TauroCA cotreatment increases WT GR exponent compared to Cort alone, while no differences are observed for GR^tetra^. Error bars represent standard error of the mean (SEM). **E-F,** Comparison of dwell time distributions between Halo-GR WT and Halo-GR^tetra^ after 1 (**E**) or 3 hours (**F**) of Cort activation, in the presence or absence of TauroCA.

A similar decrease in the dwell time of WT GR was observed when comparing 1 versus 3 hours of Cort treatment alone. In addition, TauroCA treatment does not affect GR dwell time after 3 hours, likely because of a significant reduction in the number of Cort-activated GR molecules under these conditions. We have previously shown that GR mediates transcriptional regulation through rapid receptor exchange and hormone recycling^105^. Thus, it is possible that prolonged hormone treatment promotes receptor turnover, thereby reducing dwell time independently of the bile acid. Overall, these findings show that bile acids rapidly interfere with GR chromatin binding and suggest that these molecules can modulate GR dynamics at the chromatin level.

### Taurocholic acid inhibits GR oligomerization and reduces its transcriptional activity in living cells

Current experimental evidence favors a pathway of GR activation that involves non-canonical receptor dimerization upon GC binding and nuclear import, followed by its tetramerization when bound to DNA^13,85,98,99^ (summarized in Fig. 6A). Our recently presented structure of GR-LBD from crystals with a very high solvent content revealed highly plastic tetrameric arrangements (PDB entry 9HDF)^85^. These tetramers are formed by the association of pairs of non-canonical GR dimers centered on loops L1-3 (core dimers) through their H9/L9-10/H10 surface, particularly SF-3 residues Trp712 and Tyr716 (Fig. 6A)^85^. The significant plasticity at this dimer-of-dimers interface allows different relative inter-dimer orientations, from more extended, ‘open’ to ‘closed’ arrangements. These conformations have notably different SF-3 accessible areas, and therefore different binding sites for small molecules (Fig. 6B). Demonstration that SF-3 preferentially recruits cholesterol-derived molecules such as bile acids and biophysical evidence presented above prompted us to explore their capacity to interfere with GR oligomerization in living cells. To this end, we performed quantitative fluorescence microscopy (number and brightness, N&B) assays, as previously described^99,106^. Briefly, 3134-GR^KO^ cells were transiently transfected with GFP-tagged GR and treated with 100 nM Cort, alone or in combination with 100 µM TauroCA. These cells express a tandem array of DNA binding sites for GR, the mouse mammary tumor virus (MMTV) array, to allow quantitative assessment of GR oligomeric state both in the nucleoplasm and when bound to DNA (Fig. S6C).

**Fig. 6.**
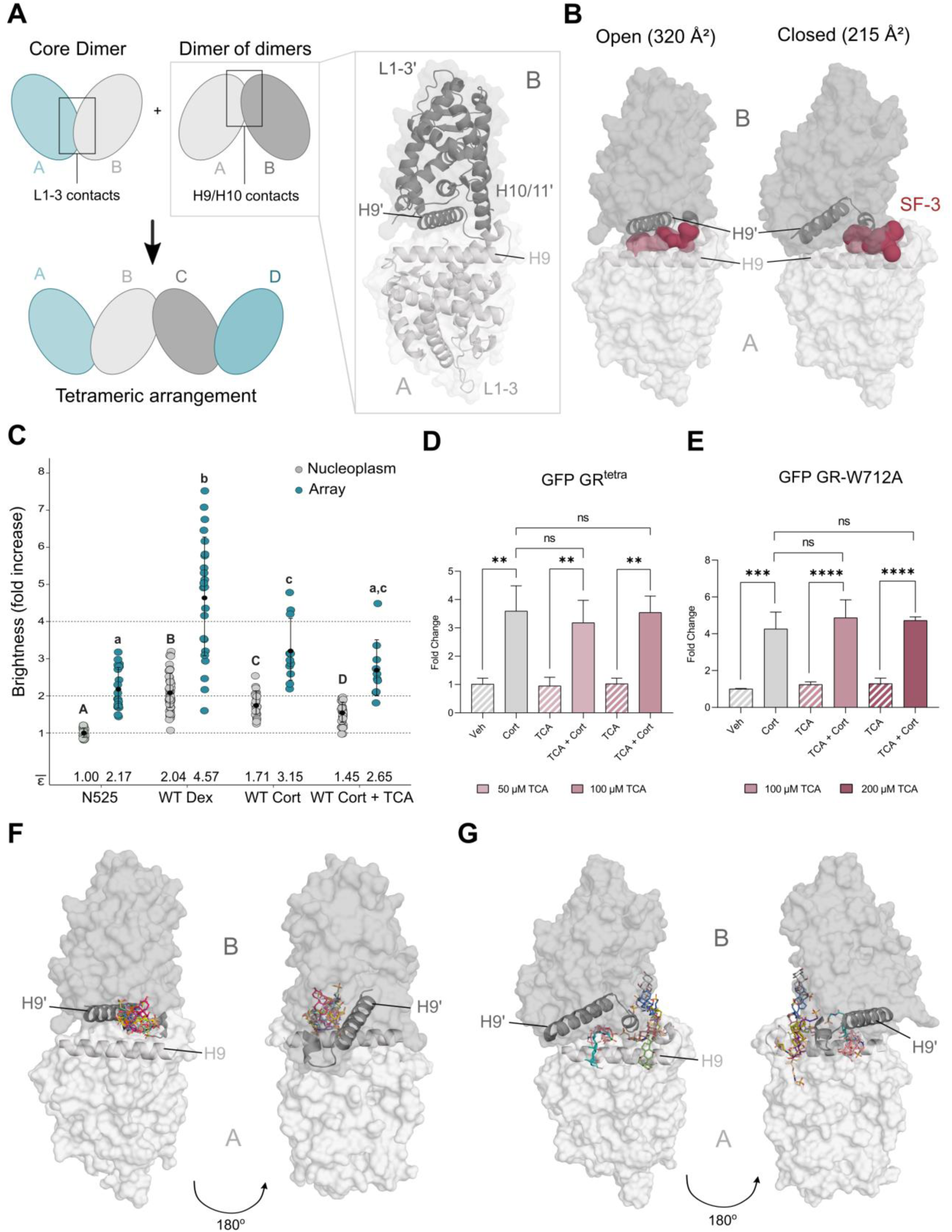
TauroCA modulates GR transcriptional activity in living cells by interfering with its self-assembly. **A**, Schematic representation of GR-LBD multimerization pathway. The core, non-canonical dimer is stabilized through L1-3 contacts, while interaction of two core dimers through the H9/H10 surface results generates a tetrameric arrangement (dimer-of-dimers). The zoomed-in structural view highlights key interface regions, including L1-3, H9 and H10/11. **B**, Open and closed conformations of GR-LBD dimer-of-dimers, indicating the variation in SF-3surface. **C**, FL GR oligomerization in the nucleus as assessed by N&B. The fold increase in molecular brightness (ε) relative to the N525* monomeric control is shown (*N* = 134 total cells with 10 < *N* < 33 between experiments). Statistical analysis was conducted independently for molecules in the nucleoplasm and at the array (capital and small letters, respectively). Different letters indicate significant differences (p < 0.05, ANOVA and Tukey test; data from four independent microscopy sessions). **D-E**, *Per1* gene induction in a stable 3134-GR^KO^ cell line expressing (**D**) GFP-mGR^tetra^ or (**E**) GFP-mGR-W718A (W712A in hGR) after 30 min of 100 nM Cort treatment, alone or in combination with TauroCA, as assessed by RT-qPCR. Mean relative expression levels for all the experiments are normalized to a housekeeping gene (β*-actin*), with error bars indicating standard deviations from three biological replicates. Statistical analysis was performed using one-way ANOVA followed by Tukey’s post-hoc test to compare group means. Significant differences (p < 0.05) are denoted by asterisks (*p < 0.05, ** p < 0.01, *** p < 0.001 and **** p < 0.0001). **G-H**, Results of ten independent docking runs of TauroCA onto the two conformations of the GR-LBD dimer-of-dimers. In the ‘open’ conformation (**G**), all runs consistently place the TauroCA molecule (colored spheres) within the extended SF-3. By contrast, in the ‘closed’ conformation (**H**) only one TauroCA molecule occupies this cavity, while the other four runs position the bile acid randomly along the dimerization interface.

Corticosterone treatment induced GR dimerization in the nucleoplasm and tetramerization at the array as expected, although to a lower extent compared to Dex-stimulated cells (Fig. 5B). When cotreated with TauroCA, Cort-induced dimerization and tetramerization of the nuclear receptor were substantially impaired with an ε decrease from 1.71 to 1.45 and 3.15 to 2.65, respectively (Fig. 6C). These results suggest that bile acids can disrupt the inter-monomer interactions needed to form and stabilize GR tetramers by binding to SF-3.

To further explore the interplay between bile acid binding and receptor multimerization, we took advantage of the constitutively tetrameric GR variant, GR^tetra^ ^98,99^. We reasoned that the multimerization behavior of this variant would be less affected by bile acid treatment compared to WT, likely because SF-3 would be at least partially occluded within GR tetramers. To verify this hypothesis, we first analyzed *Per1* expression in 3134-GR^KO^ cells expressing GFP-tagged GR^tetra^ and treated with 100 nM Cort alone or in combination with 100 µM TauroCA. As expected, no downregulation of GR function was observed for the constitutive tetrameric variant (Fig. 6D). These results are in line with the lack of effect on chromatin binding observed for GR^tetra^ after TauroCA treatment (Fig. 5C). To further explore this, WT-GR and GR^tetra^ were treated with 100 nM Cort and their transcriptional programs in the presence and absence of 100 µM TauroCA were compared. Transcriptional activity of Cort-induced GR was reduced by more than 80% after TauroCA treatment, whereas GR^tetra^ response was not affected (Fig. S6, D to F).

To verify that bile acids modulate GR tetramerization by binding to SF-3, thus interfering with inter-monomer interactions between aromatic residues at the N-terminal end of H10, we performed further experiments with the GR^W712A^ mutant (Trp718 in mouse GR). We have previously shown that this truncation does not impair GR oligomerization on its own^13^, but it would alter the SF-3 architecture and reduce bile acid binding, allowing us to target this cavity without altering central GR functions. In striking contrast to the results obtained with WT GR (Fig. 4B), *Per1* expression levels in cells expressing the GR^W712A^ mutant after 30 minutes of treatment with 100 nM Cort were not affected by the presence of 100 µM TauroCA (Fig. 6E). These findings strongly support our hypothesis that bile acids interact with GR in living cells mostly through SF-3, thus interfering with GR tetramerization and chromatin binding and therefore with its transcriptional activity.

To further explore bile acid binding modes to SF-3, we performed docking experiments of TauroCA on monomeric GR-LBD and two distinct conformations of the “dimer of dimers” generated by juxtaposition of the Trp712-centered interfaces from two monomers^85^ (Fig. S5G and Fig. 6, F and G). In the five independent docking runs on the open dimer, TauroCA was consistently positioned within the SF-3 cavity (Fig. 6F), highlighting the accessibility of this pocket to bile acids in this arrangement. In contrast, only one docking run on the closed conformation positioned TauroCA within the SF-3 cavity, while the other nine placed the molecule randomly along the dimer interface (Fig. 6G). Moreover, important clashes of the docked TauroCA molecules with residues Lys695, Trp712, and Tyr716 can be observed when superimposing molecules docked in the open arrangement on the closed conformation (Fig. S5H). Thus, the reduced size and accessibility of the SF-3 pocket in the closed dimer-of-dimers of GR-LBD would prevent effective bile acid binding.

## DISCUSSION

Our findings illuminate previously unappreciated mechanisms through which bile acids modulate glucocorticoid receptor activity. While it has been assumed that bile acids can compete with GCs for the internal ligand-binding pocket of this nuclear receptor, our current structural and docking evidence indicates that this regulatory role is actually driven by the occupancy of well-defined cavities on the surface of the ligand-binding domain. Solvent-exposed surfaces enriched in hydrophobic/aromatic residues are primary binding sites for chaperones and cochaperones, but in isolated proteins, they recruit small molecules with detergent-like properties instead^18^. In line with this observation, a thorough bioinformatics analysis of the LBD surface and previous structural evidence reveals two major GR-LBD cavities as important bile acid targets, for which we have coined the terms SF-3 and E sites. The E site is spanned by residues of loops L1-3 / L6-7 and H11, while the SF-3 cavity is delimited by residues in helices H9/10 and the connecting loop, most notably the conserved aromatic residues at the N-terminus of H10, Trp712 and Phe715 (Fig. 1, D to H). Noteworthy, these cavities partially overlap with two major inter-monomer interfaces at the top and back of the GR-LBD module in the standard orientation (Fig S1B, C)^13,85^.

Our current results indicate that both short- and long-term effects of bile acids on GR transcriptional activity are largely a consequence of the competition between bile acid binding and receptor self-assembly and underscore the importance of SF-3 in this regard. (1) N&B results show a direct impact of TauroCA binding on receptor di- and tetramerization (Fig. 6C), similar to what we have previously observed for SF-3 mutants, GR^W712E^ and GR^W712S/Y716S^ ^85^; (2) Chromatin binding and transcriptional activity of the WT receptor, but not of the constitutively tetrameric variant (GR^tetra^), are altered by short-term bile acids treatment (Figs. 5C and S5F); (3) Bile acids are unable to downregulate GR transcriptional activity in cells stimulated with 100 nM Dex or 600 nM Cort (Fig. S2, D and E) suggesting that sustained GR activation promotes the formation of stable tetramers leading to occluded bile acid binding sites; (4) Long-term bile acid treatment protects GR from proteasomal-driven degradation and reverts corticosterone-induced repression of *GR* expression.

The most common arrangements of GR-LBD dimers-of-dimers in our recently reported structure of multimeric GR-LBD positions the SF-3 pockets of two monomers in close proximity, thus generating a large, solvent-protected area enriched in aromatic residues^85^. This arrangement has been observed in several additional crystal structures, some of which feature a detergent molecule occupying the SF-3 cavity of two neighboring monomers^13^. These properties are characteristic of so-called multichain binding sites, i.e., binding sites for small-molecule ligands formed at the interface between two or more monomers^107,108^. These studies demonstrated that residues within multichain binding-sites are better conserved overall than those involved in forming single-chain binding sites. Indeed, the residues that span SF-3 are highly conserved in GR across species, ranging from fishes to humans, most notably the triplet of aromatic residues in H10 (Fig. 1D). Our docking experiments revealed that bile acid molecules would bind across two neighboring SF-3 sites in the more open arrangement of Trp712-centered dimers-of-dimers, as observed in crystal structures (Fig. 1E). However, in the ‘closed’ conformation the bile acid moiety would clash with residues of the second LBD monomer (Fig. S5H). Thus, bile acid binding would interfere with the ratcheting of GR dimers-of-dimers hindering their ability to transition between different relative conformations. This interference might impair optimal binding to chromatin and/or dynamics of the transcription machinery, ultimately leading to the observed decrease in transcriptional output. Altogether, our results suggest that bile acid binding to the GR tetramer is highly dependent on the specific conformation of the Trp712 interface, with the accessibility of the SF-3 cavity playing a crucial role in GR modulation.

GR relies on a complex interplay between chaperones (Hsp70/Hsp90), cochaperones (p23) and immunophilins (FKBP51, FKBP52) to adopt its mature, transcriptionally active conformation^28,109,110^. Notably, SF-3 overlaps with the binding site for cochaperone p23 (Fig. S1E)^28^. We have observed that GR translocation to the cell nucleus is not impaired by bile acids, but interference with GR-nuclear chaperone interactions might affect the proper cycling of the receptor on chromatin^111^, further contributing to the reduction in transcriptional activity by potentially impairing its dynamic engagement with target gene loci. GR appears to depend on the nuclear chaperone function as part of the nuclear quality control system (involving an interplay with the ubiquitin-proteasome system) for its proper function as a transcription factor *in vivo* ^105^. Of note, our current results after long-term treatment indicate that bile acids not only interfere with the proteasomal-driven GR degradation but also inhibit a hormone-induced inhibitory feedback loop^92–94^. A schematic summary of the manifold effects of bile acids on GR function *in vivo* is shown in Figure 7.

**Fig. 7.**
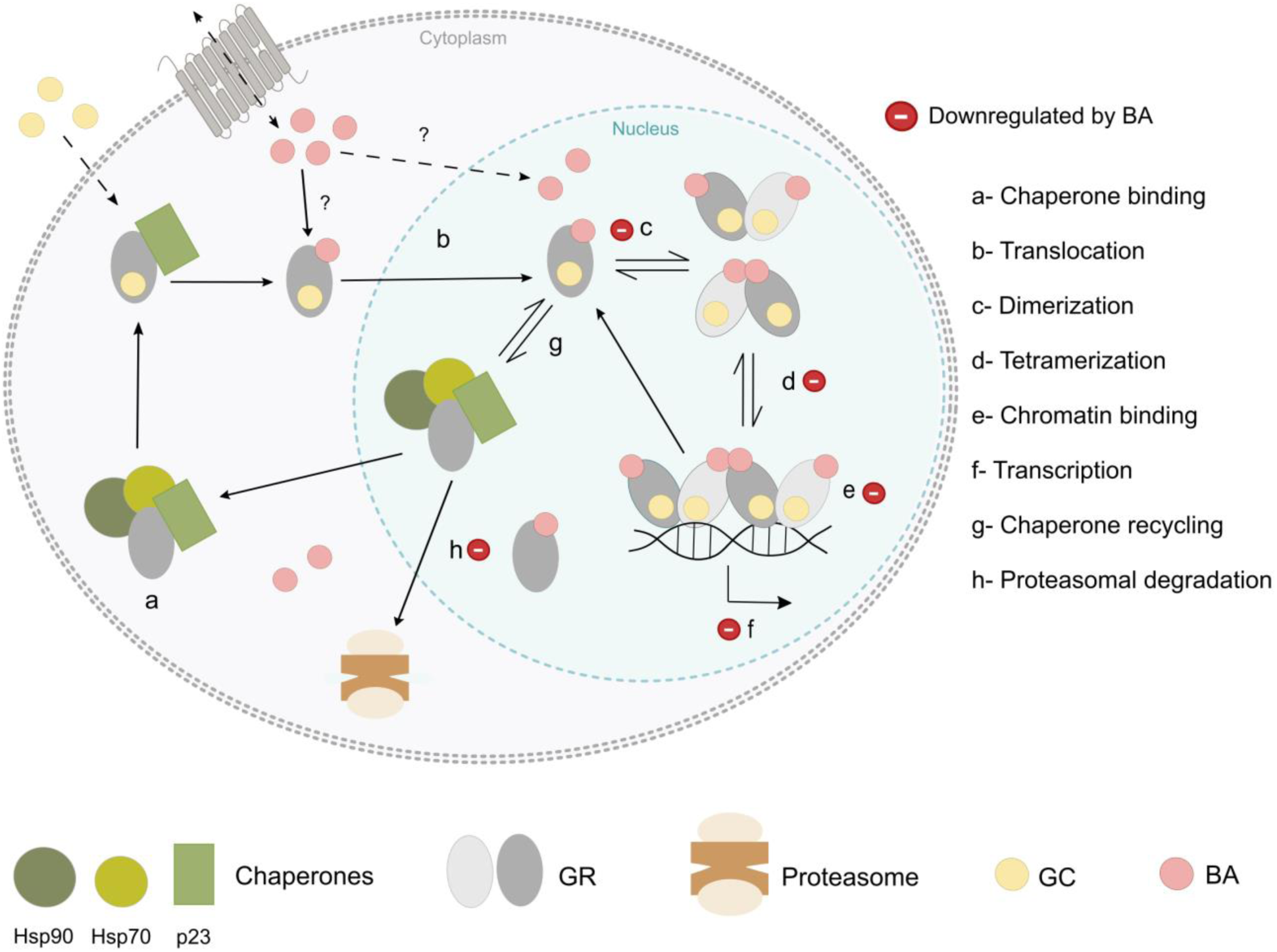
Mechanisms of bile acid-mediated regulation of GR function. Schematic representation of bile acids effects on the life cycle of the glucocorticoid receptor. Bile acids can then either diffuse into the nucleus or be actively transported bound to other proteins, including GR^117^. In the cytoplasm, inactive GR is stabilized by a chaperone complex comprising Hsp90, Hsp70, and p23 (a). Upon glucocorticoid binding (GC, yellow circles), GR undergoes conformational changes and translocates into the nucleus (b). Inside the nucleus, GR forms dimers (c) which then associate into tetramers (d) upon chromatin binding (e). This leads to transcriptional activation of target genes (f). In the nucleus, GR cycles between GC-bound (active) and GC-free conformations, in a process that is also facilitated by the chaperone machinery (g). Finally, GR can be either exported to the cytoplasm or degraded by the proteasome (h). Current experimental data shows that bile acids (BA, pink circles) significantly downregulate GR function in the short- and long-term by interfering with several of these key steps, including receptor turnover, oligomerization, and chromatin binding (highlighted with a negative icon). By interfering with GR structure and function at multiple levels, bile acids significantly impair GR-mediated gene regulation, reducing its transcriptional activation of target genes. This is in addition to the inhibition of Cort-initiated negative feedback loop.

The dynamic equilibrium between GR monomers and oligomers depends on the potency of the LBP-bound glucocorticoid. Our N&B analysis, which allows for the *in situ* quantification of GR oligomerization states in living cells indicates that the less potent natural GC, corticosterone, has a lower ability to shift the equilibrium towards multimers compared to Dex. This balance is significantly displaced towards oligomers for GR^tetra^, and consequently, less dynamic. Thus, bile acids have higher probability of interacting with WT-GR compared to GR^tetra^, as the SF-3 cavity is fully accessible in GR homodimers. The impact of bile acids binding to SF-3 on both dimerization and tetramerization is also in line with allosteric rearrangements that extend to the opposite pole of the domain, including the dimerization site centered on loop L1-3, as previously observed by us and others^112,113^. Bioinformatic analyses revealed that the SF-3 cavity is unique for GR, and even in the closely related MR, no evidence for small-molecule binding to the topologically equivalent area has been reported to the best of our knowledge. Inspection of deposited crystal structures of MR-LBD reveals a tight packing of monomers that leaves no space for cholesterol-derived molecules close to the conserved Trp712/Phe715/Tyr716 triad (human GR numbering; see e.g. the structures of MR-LBD bound to Cort, aldosterone or progesterone, PDB entries 2AAX, 2AA2 and 2AA5, respectively^114^). These observations suggest that bile acids could selectively modulate GR activity without affecting MR, highlighting a potential for targeted regulation without cross-reactivity.

On the other hand, binding to the other major cavity on the GR-LBD surface, the E site is likely to have direct effects by interfering with hormone trafficking in and out of the LBP, in addition to impacting receptor multimerization. Direct physical interactions between small molecules bound at the E site and inside the LBP have been reported. For instance, a HOG molecule in PDB 3K22 makes VdW contacts with the bound GC (Fig. S1D)^80^. Direct contacts between lithocholic acid that occupies the E site and the LBP-bound 4-hydroxy-tamoxifen have also been reported for estrogen receptor-related γ (ERRγ) (PDB 1S9Q^115^; 7R62^116^).

We stress that bile acid concentrations used in our experiments in living cells are within the range of those observed *in vivo*, mainly in pathological conditions such as cholestasis or liver dysfunction^35,45,46,50,51,62,118,119^. In our hands, TauroCA showed a slightly higher impact on GR transcriptional activity than bile acids such as LCA and UDCA. These functional differences are most likely related to the presence of a longer, acidic taurine tail, which is capable of extending towards the Arg690 side chain, engaging in salt bridge formation with its guanidinium group. Future investigations should determine whether other secondary bile acids bind with even higher affinity to GR-LBD, potentially exerting a greater impact on the transcriptional activity of the receptor. Given the vast diversity of bile acids^119–121^, it is tempting to speculate that direct and specific links connect the gut microbiota with major physiological processes through direct physical interactions with nuclear receptors^122^.

This bile acid-mediated modulation of steroid receptors through binding to a solvent-exposed cavity distant to the internal hormone-binding pocket is conceptually similar to exosites identified in regulatory serine proteases^123^. This is in strong contrast to other previously characterized BARs, FXR and VDR, and presumably other members of the NR1H and NR1I subfamilies, in which bile acids occupy the internal LBP^58^. Interestingly, similar to the regulatory serine proteases, dual binding modes have been previously reported for bile acids. Specifically, binding of a second LCA molecule to the VDR surface was shown to be critical for LCA agonism^17^. Current evidence indicates that bile acids play a central role by modulating the intertwined relationship between the two groups of nuclear receptors, especially between GR and FXR^124^. In this regard, LCA has been recently shown to mediate the effect of calorie restriction by activating the master regulator of metabolic homeostasis, AMPK^125,126^, which modifies transcription of GC-responsive genes in a tissue- and promoter-specific manner upon phosphorylation of GR’s N-terminal domain^127^.

Furthermore, biologically relevant bile acid derivatives with high affinity for the LBP of oxosteroid receptors can be generated by specific modifications of the cholesterol core. This has been recently demonstrated with the discovery of the LCA derivative, 3-oxo-D^4,6^-LCA, which potently and selectively binds AR and antagonizes its transcriptional activity, thus limiting prostate cancer growth and metastasis^122^. This finding highlights the potential for designing bile acid-based modulators of steroid receptor activity. Docking experiments indicate that the LCA isomer would bind in the opposite orientation as endogenous AR ligands such as dihydrotestosterone. A few conservative replacements in the E site between AR and other oxosteroid receptors are likely responsible for this unique specificity of 3-oxo-D^4,6^-LCA. For instance, the residue topologically equivalent to Tyr735 in the GR is a phenylalanine in the AR. This residue is important for GR transactivation, which is reduced by 30% in the Tyr735Phe mutant^128^. Our docking experiments of 3-oxo-D^4,6^-LCA on GR-LBP show that it behaves similarly to TauroCA (Fig. S2H and I), binding mainly outside the internal pocket.

Despite the extraordinary success of drugs that target the internal ligand-binding pockets of several nuclear receptors (e.g., antiandrogens for prostate cancer treatment, synthetic glucocorticoids for treating asthma and other inflammatory conditions), the rapid emergence of resistance and the significant side effects of these LBP-directed drugs has fueled the search for new NR antagonists with alternative mechanisms of action^18^. Solvent-exposed binding pockets on the surface of the LBD modules are attractive alternative targets for pharmacological intervention.

Our current results offer a compelling rationale for developing new GCs with reduced side effects by exploiting the biochemical and biophysical properties of bile acids, including their biocompatibility. This approach opens avenues for designing more selective and physiologically compatible modulators of GR activity. Previous observation that a synthetic GC, tailored for binding to the LBP, also binds tightly to the SF-3 pocket of GR-LBD^84^ and the proven pharmacological relevance of TauroCA in a model of ataxia^75^ demonstrate both the feasibility and the potential clinical relevance of this approach.

## MATERIAL AND METHODS

### Bioinformatics analyses

PDB entries 1M2Z, 7YXO and 4UDD were used as templates to run Fpocket, PockDrug and CavitySpace algorithms, respectively, using the standard parameters^76–78^. Analog screenings were performed using fingerprint (FP) similarity and substructure searches in the Zinc15 ^129^ and DrugBank databases^130^. The FP search with the Erlwood FP similarity node used Morgan Fingerprints with radii of 2 and 1024 bits. The substructure search was based on SMARTS patterns. For the FP similarity search, we applied a similarity cut-off of 0.6. AutoDock Vina^131^ and Glide SP docking algorithms (www.schrodinger.com/platform/products/glide/) were used to identify potential docking conformations and affinities of selected compounds (Table S4) with monomeric and dimeric GR-LBD structures using PDB 4UDD and 9HDF. Prior to docking, receptor and ligand molecules were parametrized by adding hydrogen atoms, removing water molecules, and computing partial atomic charges. For AutoDock experiments, an exhaustiveness parameter of 20 was applied to ensure thorough conformational sampling. The number of docking runs and the energy range values were set to 10 and 5 kcal/mol, respectively, to promote diversity in binding poses and capture a wide range of energetically favorable interactions for each ligand.

#### AlphaFold3 structure predictions

Model generation with AlphaFold3 (https://alphafoldserver.com/) was performed using 20 randomly generated seeds. GR-LBD sequence was used along with different combinations of two ligands: corticosterone (C0R) and taurocholic acid (TCH), employing ccd codes to ensure the proper chirality of the compounds during the AF3 execution. The ranking scores obtained for all models were consistently over 0.8. <u>Calculation of binding energies:</u> Binding energies were calculated using the pyDock energy scoring function, which uses an adapted version of the pyDock scoring function, originally developed for protein–protein interactions, to evaluate protein-ligand energetic profiles, including the electrostatic, van der Waals, and desolvation terms. This scoring function was successfully used in the 7^th^ CAPRI edition^132^. The structural data of the ligands was processed with AmberTools23 (www.ambermd.org/) to generate the required ligand parameters. First, the net charges of the ligands were extracted from the PDB data and used for assigning partial charges. The AM1-BCC method^133^ was then used to assign partial charges to the ligands based on the net charges. Once the ligand parameters were established, pyDock was used to calculate the binding energy, allowing us to compare the energetic profiles across the different models and binding sites. Docking results were visualized and analyzed with PyMOL (www.pymol.org/), which was also used to prepare all structure figures.

### Surface plasmon resonance (SPR) assays

SPR analyses were performed at 25 °C in a BIAcore T200 instrument (GE Healthcare). Highly purified, Dex-bound recombinant WT ancGR2-LBD was diluted in 10 mM sodium acetate, pH 5.0, and directly immobilized on CM5 chips (GE Healthcare) by amine coupling at a density of 100 RUs for protein-protein interaction experiments, or 7000 RUs for protein-small molecule interaction experiments. As a reference, one of the channels was also amine-activated and blocked in the absence of protein. The running buffer was 20 mM HEPES, pH 7.2, 200 mM NaCl, 1 mM DTT, 3.3% glycerol, 0.05% Tween 20. For protein-protein interaction experiments, a constant concentration of 1 mM CHAPS / BOG or 40 µM of LCA was kept throughout the experiments. Sensorgrams were analyzed with the BIAcore T200 Evaluation software 3.0 using the results of experiments conducted with running buffer alone as baseline and fitted according to the Langmuir 1:1 and multisite models.

### Cell lines and plasmids

GFP-tagged variants of mouse GR (WT, P481R and W718A) used for RNAseq and N&B experiments were generated with the QuikChange II XL Site-Directed Mutagenesis Kit (Stratagene) and stably integrated into the GT-Rosa26 locus via CRISPR/Cas9 homology-directed repair with puromycin selection and fluorescent-activated cell sorting.

pHalo-H2B used for SMT experiments was generated by polymerase chain reaction (PCR) amplification of the H2B coding region from an H2B-GFP template and cloned into a pFC14A backbone (Promega, Madison, WI) to fuse the HaloTag to the C terminus of H2B^134^. 3134 GR KO cells expressing GFP-mGR were used to generate the Halo-mGR expressing cell line by replacing the tag using site-directed mutagenesis PCR.

Mammary adenocarcinoma 3134-derived GR knock-out (GR-KO) cells expressing GFP-tagged mouse GR variants were grown in Dulbecco’s modified Eagle’s medium (DMEM, Invitrogen) supplemented with 5 µg/ml tetracycline (Sigma), 10% fetal bovine serum (FBS, Sigma), sodium pyruvate, non-essential amino acids, and 2 mM L-glutamine. Cells were maintained in a humidifier at 37 °C. The FBS-supplemented medium was replaced with medium supplemented with charcoal/dextran-treated FBS to remove GCs from cells for 24 h prior to hormone treatment.

The Halo-GR cell line was generated by CRISPR/Cas9 targeted insertion of a Halo tag into the N terminus of the GR gene at its endogenous locus in the mammary adenocarcinoma 3134-derived cells. Treatment with IPTG induces expression of shNIPBL in these parental cells. However, under the normal culturing conditions used in this study they have normal NIPBL levels and GR-regulated gene responses^104^.

### Translocation assays

Prior to imaging, cells expressing GFP-GR were plated in 384-well plates (Matrical) at a density of 10,000 cells per well in DMEM medium containing 10% charcoal stripped serum (Hyclone, Logan, UT). Cells were activated with 100 nM Cort and treated with various concentrations of LCA or TauroCA for 2 h. After treatment, cells were fixed with 4% paraformaldehyde in PBS for 10 min and washed three times with PBS. Cells were further stained with DAPI (4′,6-diamidino-2-phenylindole, SC-3598, ChemCruz) dissolved in PBS (5 µg/ml) for 10 min, washed three times with PBS, and either imaged immediately or kept in PBS at 4 °C for later imaging.

A Yokogawa CV7000S high-throughput dual spinning disk confocal microscope was used for fully automated collection of images. Images were acquired with a 40X Olympus Plan ApoChromat air objective (NA 0.9) and two sCMOS cameras (2560 × 2160 pixels) using camera binning of 2×2 (Pixel size 325 nm). A single imaging plane was used and the 2×2 binning allowed us to reduce the size of the files. Samples were imaged using 405 nm, 488 nm, and 561 nm excitation lasers and a fixed 405/488/561/640 nm dichroic mirror on the excitation side. On the emission side, the emitted light was split by a fixed 561 nm emission dichroic mirror. Specifically, emitted light with wavelengths below 561 nm was reflected by the emission dichroic mirror at a 90-degree angle to the camera switchable 445/45 or 525/50 nm emission bandpass filters mounted on a filter wheel (Camera #1). Emitted light with wavelengths above 561 nm passed through the mirror and to the other camera with switchable 600/37 or 676/29 nm emission bandpass filters mounted on another filter wheel (Camera #2). Camera #1 and Camera #2 were positioned perpendicular to each other. DAPI (Ex. 405 nm—Em 445/45 nm) and GFP signals (Ex. 488 nm— Em. 525/50 nm) were always acquired in separate exposures. Additional images were taken on a PerkinElmer Opera QEHS High-Content Screening platform (Waltham, MA). This system employed 40× water-immersion objective lens, a laser illuminated dual Nipkow spinning disk, and cooled charge-coupled device cameras to digitally capture high-resolution confocal fluorescence micrographs (323 nm pixel size with 2×2 camera pixel binning). On average, eight fields per well were imaged using a single imaging plane.

### Single molecule tracking (SMT)

#### Microscopy

All SMT experiments were performed on a custom-built, three-color highly inclined laminated optical (HILO) sheet in the LRBGE Optical Microscopy Core at the National Cancer Institute (NCI), National Institutes of Health (NIH). This system contains a 100x 1.49NA TIRF objective (Olympus Scientific Solutions, Waltham, MA), three lasers (488, 561 and 647 nm, Coherent, Inc., Santa Clara, CA), and a stage-top incubator with CO_2_ control (Okolab, Pozzuoli NA). The emitted light passes through a quad-band dichroic (ZT405/488/561/647, Chroma, Bellows Falls, VT) and two long-pass filters (T588lpxr, T660lpxr), and is finally separated by emission filters (525/50, 609/58, 736/128; Semrock) before being detected simultaneously by three EMCCD cameras (Photometrics, Tucson, AZ). The pixel size for this system is 144 nm. For each cell, 600 frames were collected with 10 ms exposures and 200 ms intervals with a laser power of 1.3 mW at the objective.

<u>Tracking:</u> Tracking was performed using TrackRecord. Each movie was filtered with a band-pass filter and the nucleus was identified from a temporal projection of the movie. Particles detection was performed using an intensity threshold where >95% of detected particles have a signal-to-noise ratio ≥ 1.5. Particles were connected to form tracks using a nearest-neighbor algorithm with the following parameters: maximum jump = 4 pixels (576 nm), gaps to close = 1 frame, shortest track = 2 frames.

#### Dwell time calculation and power-law fit

Histone H2B was used to calculate the parameters of bound molecules as described in^100^. Following the published procedure herein, the photobleaching rate k_PB_ was calculated by fitting the H2B survival distribution to a triple-exponential function. Dwell time distributions of bound GR WT and GR^tetra^ molecules were corrected for photobleaching by dividing the respective raw survival distributions by the exponential photobleaching component:

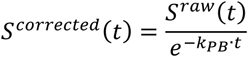

The resulting survival distribution was then fit to double-exponential and power-law models, and a Bayesian information criterion was used to determine the best model fit as described^100^.

#### N&B experiments

Quantitative fluorescence microscopy images were taken at the CCR, LRBGE Optical Microscopy Core facility (Bethesda, MD) using LSM 780 laser scanning microscopes (Carl Zeiss, Inc.), equipped with an environmental chamber. Cells were excited with a multi-line Argon laser tuned at 488 nm and imaged for 20-120 min after hormone addition using a 63× oil immersion objective (NA = 1.4). Fluorescence was detected with a GaAsP detector in photon-counting mode, using 491-589 nm filtering, N&B measurements were done as previously described^13,99^. For each studied cell, a single-plane stack of 120 images (256 × 256 pixels) was taken with a pixel size of 80 nm and pixel dwell times of 6.27 µs in the LSM 780 scope. In all stacks, we discarded the first 5 images to reduce the effect of photobleaching. The frame time under these conditions is 0.97 s, which guarantees independent sampling of molecules according to previously reported fluorescence correlation spectroscopy (FCS) measurements^135^. Each stack was further analyzed using the N&B routine of the SimFCS 2.0 software (Global Dynamics), in which the average fluorescence intensity () and its variance (σ2) at each pixel of an image are determined from the intensity values obtained at the given pixel along the image stack. The apparent brightness (B) is calculated as the ratio of σ2 to while the apparent number of moving particles (N) corresponds to the ratio of to B^136^. We used the SimFCS 2.0 software to classify pixels as occurring in the nucleus or the MMTV array according to their intensity values^99^.

Cells were selected for analysis following these criteria: (i) in the case of stimulated cells, an accumulation of signal at the array must be visible, (ii) the average apparent number of molecules (N) in the nuclear compartment must have a range of 3–30 units in all cases, (iii) no saturation of the detector at any pixel (N < 60), and (iv) bleaching cannot exceed 5–10%. In previous work it was demonstrated that B is equal to the real brightness ε of the particles plus one^136^. Therefore, ε at every pixel of the images can be easily extracted from B values. Importantly, this analysis only provides information regarding the moving or fluctuating fluorescent molecules since fixed molecules (relative to our frame time) will give B values equal to 1. The oligomerization state is obtained by comparing the fluorescence of full-length GR to that of a constitutively monomeric GR variant, N525*. The experiments were independently repeated at least two times for each treatment/condition.

#### Expression analysis of GR target genes by RT-qPCR

For time-course experiments and qPCR, GFP-GR and endogenous GR expressing cells were left untreated or treated with 100 nM of Cort for 0.5, 1, 2 or 3 h prior RNA isolation, in the presence or absence of bile acids at different concentrations. RNA was isolated using the PureLink RNA kit (Thermo #12183018 A; Thermo Fisher Scientific) and cDNA reactions were performed with the NEB Protoscript II kit (#E6560S) following the manufacturers’ instructions. RT-qPCRs were performed with iQ SYBR Green Supermix (Biorad # 1708880) in a Biorad CFX96 machine with duplicate wells for each sample. The following primers were used to amplify *Per1* (5’-CTTCTGGCAATGGCAAGGACTC-3’; 5’-CAGCATCATGCCATCATACACAC-3’), *Sgk1* (5’-GAAACAGAGAAGGATGGGCCTGAAC-3’; 5’-GATCTCAGCTCCAGCACCACCAC-3’), *Arl4d* (5’-AAAAGGAATATTTAGAAGGGTAGGGTGC-3’; 5’-GCCATTTCAGTCAA-GTGGTTCCC-3’), *Tsc22d3* (5’-ACATGATGGTGGCATGAAGA-3’; 5’-TCTTCTCAAG-CAGCTCACGA-3’), *Nr3c1* (5’-GACTTCCTTGGGGGCTATGAA-3’; 5’-AACAGCAAAGTGCTGACTTAC-3’), (*β-actin* (5’-GCTGGAAAAGAGCCTCAGGGC-3’; 5’-CGCATCCTC-TTCCTCCCTGGAG-3’) and *18-S* (5’-GCCCGAAGCGTTTACTTTGA-3’; 5’-TCCATTATTCCTAGCTGCGGTATC-3’) genes. Results were normalized to either *β-actin* or *18-S* expression levels, and each sample was quantified in triplicate. GraphPad Prism 8.0 software was used to perform statistical analyses using one-way ANOVA followed by Tukey’s post-hoc test to compare group means.

#### Total RNA collection and sequencing

RNA was isolated using the PureLink RNA kit following the manufacturer’s instructions. We collected three biological replicates of each condition and assessed sample quality using the Agilent Bioanalyzer. Strand-specific sequencing libraries were generated from rRNA-depleted (Illumina RS-122-2301) total-RNA samples, using Illumina Stranded Total RNA (Illumina20020596) according to the manufacturer’s instructions. Raw reads were demultiplexed into Fastq format allowing up to one mismatch using Illumina Bcl2fastq v2.17. Reads of the samples were trimmed for adapters and low-quality bases using Cutadapt 1.18. RNA-seq alignment to the mouse mm10 genome was performed with STAR^137^. DESEQ2^138^ was used to normalize the data by read depth, identify differentially expressed genes for each form of GR (Cort/vehicle), and calculate log2 fold changes (FC) and false discovery rates (FDR) for each gene.

## Abbreviations

AF: Activation function
AR: Androgen receptor
BF-3: Binding function-3
BOG octyl: beta-D-glucopyranoside
CA: Cholic acid
CDCA: Chenodeoxycholic acid
CHAPS: 3-[(3-cholamidopropyl) dimethylammonio]-1-propanesulfonate
Cort: Corticosterone
DCA: Deoxycholic acid
Dex: Dexamethasone
ER: Estrogen receptor
E site: Entry/exit site
FXR: Farnesoid X receptor
GC: Glucocorticoid
GR: Glucocorticoid receptor
LBD: Ligand-binding domain
LBP: Ligand-binding pocket
LCA: Lithocholic acid
MR: Mineralocorticoid receptor
N&B: Number and brightness
PDB: Protein Data Bank
PR: Progesterone receptor
RNA-seq: RNA sequencing
SF-3: Sensor function-3
SMT: Single-molecule tracking
SPR: Surface plasmon resonance
TauroCA: Taurocholic acid TauroUDCA
Tauroursodeoxycholic acid: UDCA Ursodeoxycholic acid
VdW: Van der Waals
WT: Wild type

## Author contributions

Conceptualization: AJP, LV, IA, JFR, PFP, GLH, EEP

Methodology: AJP, TAJ, KW, AAM, IMN, ALL, UE, MS, MA, DMP, DAS

Investigation: AJP, TAJ, KW, AAM, IMN, ALL, UE, MS, MA, DMP, DAS

Visualization: AJP, TAJ, KW, AAM, IMN, IG, ALL, UE, PFP

Funding acquisition: AJP, AAM, JFR, GLH, EEP

Project administration: PFP, GLH, EEP

Supervision: PFP, GLH, EEP

Writing – original draft: AJP, PFP, EEP

## Data and materials availability

The Gene Expression profiles are deposited in the Gene Expression Omnibus (GEO) database and the code assigned is GSE287351. MATLAB-based tracking software is publicly available at https://doi.org/10.5281/zenodo.7558712. All data reported in this paper will be shared by the lead contact upon request. Requests for further information or reagents should be directed to and will be fulfilled by the lead contact, Eva Estébanez-Perpiñá. No generative AI or AI-assisted technologies were used at any stage.

