## Supplementary material for "Bile acids target an exposed cavity in the glucocorticoid receptor modulating receptor self-assembly, chromatin binding and transcriptional activity": SI

**The PDF file includes:**

Figs. S1 to S6

Tables S1 to S4

Legend for Movie S1

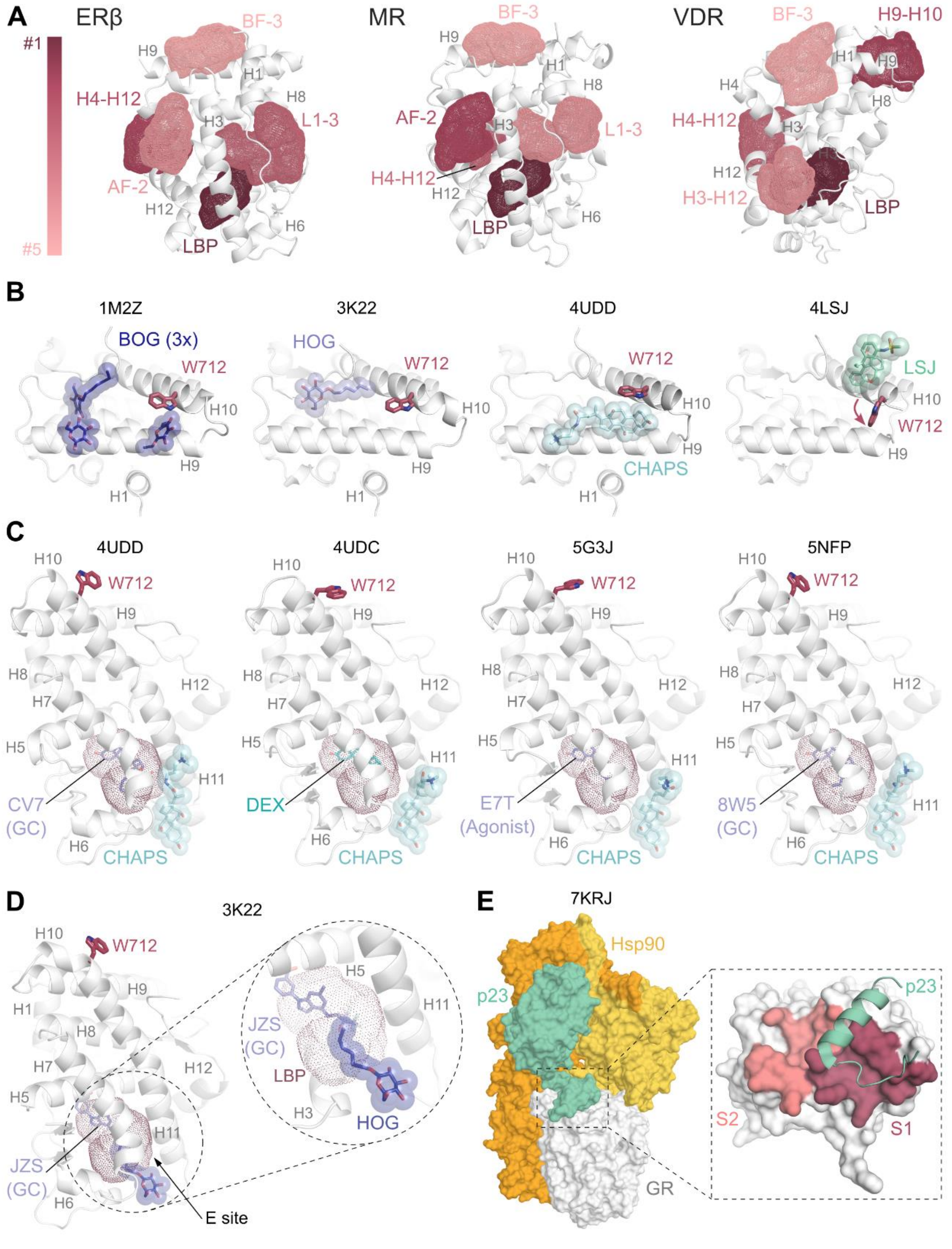


**Fig. S1. Detailed analysis of small-molecule binding sites on the GR-LBD surface.**

**A**, Representation of the top five cavities in the ligand-binding domain of steroid receptors ERβ and MR as compared to VDR. Major secondary structure elements are labeled. Pockets are shown color-coded based on their druggability scores, as in Fig. 1A. **B**, Cartoon of the GR-LBD surface, highlighting small molecules co-crystallized in the indicated PDB entries. The indole ring of Trp712 is shown in all panels; note that it swings to accommodate different compounds in the S site. **C**, A second binding site for small molecules, most notably CHAPS, has been identified in several crystal structures close to the LBP and has been termed entry (E) site. Molecules occupying the two cavities are shown and labeled. **D**, Structure of GR-LBD highlighting the HOG molecule bound in the E site. Note that the tail of the HOG molecule protrudes into the LBP and contacts the bound glucocorticoid. **E**, The co-chaperone, p23, interacts with the S site of GR-LBD. Three-dimensional structure of the GR-LBD complex with the chaperone, Hsp90, and the co-chaperone, p23. The inset shows the C-terminal helix from p23 bound across the S site. See the main text for details and references.

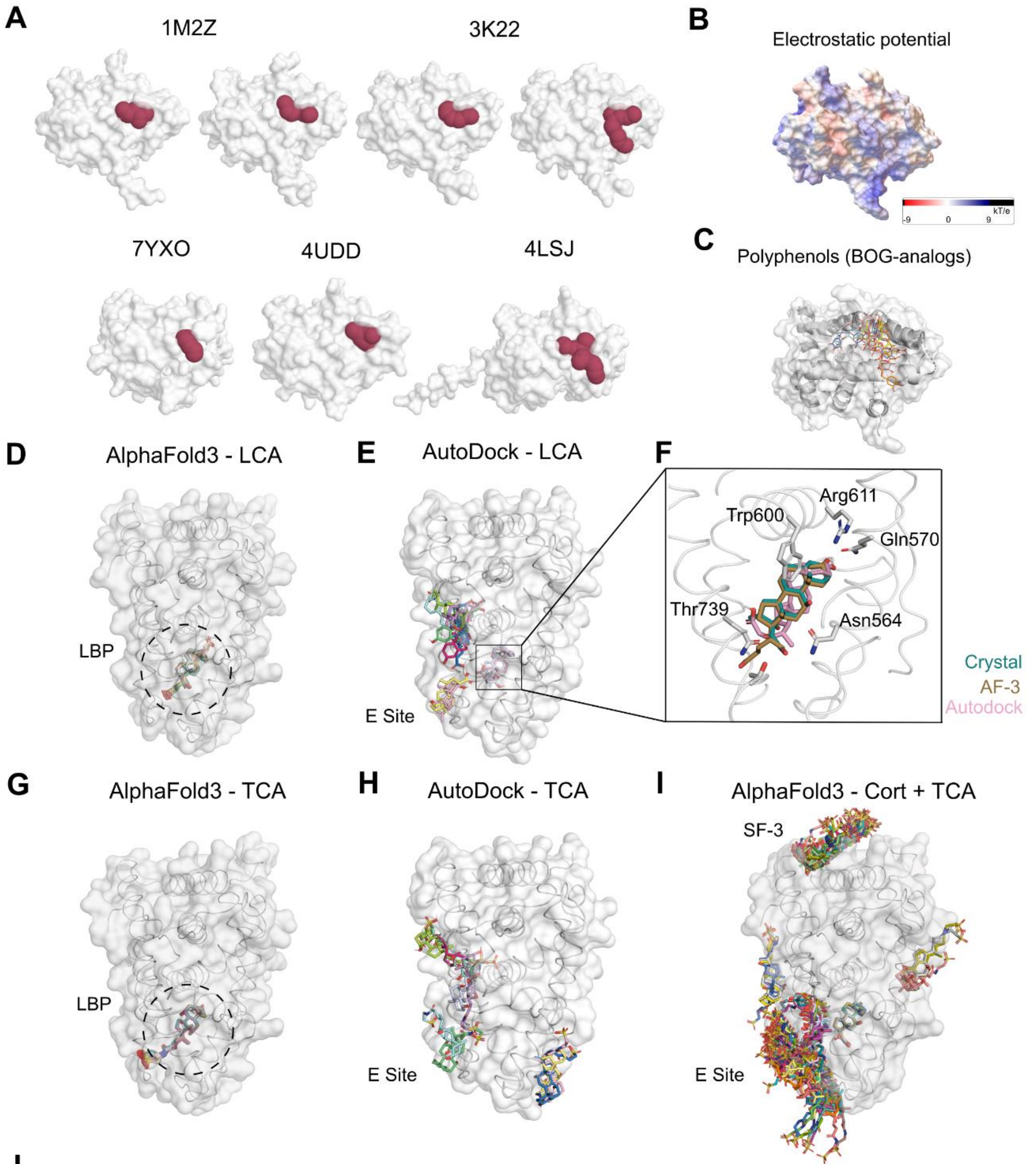


**J**

| | AlphaFold binding energies (pyDock) | | | |
|---|---|---|---|---|
| | Electrostatic | Desolvation | VdW | Total |
| **LBP** | -3.36 ± 0.18 | -6.33 ± 0.15 | -46.20 ± 1.01 | -14.30 ± 0.25 |
| **SF-3** | -0.17 ± 0.91 | -9.80 ± 1.14 | -20.78 ± 3.38 | -12.05 ± 2.09 |
| **E Site** | -0.49 ± 2.1 | -6.50 ± 1.25 | -28.71 ± 3.46 | -9.86 ± 2.08 |

**Fig. S2. Computational analysis of small-molecule binding to SF-3 as compared to the LBP.**

**A**, Surface representation of GR-LBD monomers featuring surface-bound small molecules in deposited crystal structures. The SF-3 cavity identified by Fpocket is shown with red spheres. Note that structural variations influence pocket accessibility and druggability. **B**, Representation of the electrostatic surface potential (Ψ) of GR-LBD calculated using AutoDock Vina. Negatively charged areas (Ψ ≤ -9 kT/e) are colored red, while blue indicates positively charged areas (Ψ ≥ +9 kT/e). Neutral regions are shown in white. **C**, Top predicted SF-3 binders identified in a screening of BOG-related compounds. **D**, LCA binding to human GR-LBD as predicted by AlphaFold3. The 100 top hits are shown; note that the algorithm consistently places the bile acid inside the LBP. **E**, Docking poses generated from 10 independent runs of LCA docking to GR using AutoDock. Note that in most cases the molecule binds in different conformations around the hormone-binding pocket, and only two poses dock inside the LBP. **F**, Closeup of the LBP showing the experimentally observed Dex molecule (PDB 7YXC) superimposed with the best AlphaFold3 model and the best docking pose for LCA. **G**, TauroCA (TCA) binding to human GR-LBD as predicted by AlphaFold3. The 100 top solutions are shown. Note that although the algorithm consistently places the bile acid inside the LBP, the tail of the compound remains outside the pocket. **H**, Docking poses generated from 10 independent runs of TCA docking into the LBP of GR generated with AutoDock. Note the diversity of binding conformations around the hormone pocket, with no poses docking inside the LBP. **I**, Superimposition of the 100 top AlphaFold3 models binding of one Cort and two TCA molecules to human GR-LBD. Cort is consistently predicted to bind inside the LBP, whereas TCA molecules explore different cavities of the GR-LBD surface, mostly the SF-3 and the E sites. **J**, Binding energies of the top 10 AlphaFold3 models featuring TCA molecules bound to both SF-3 and the E-site.

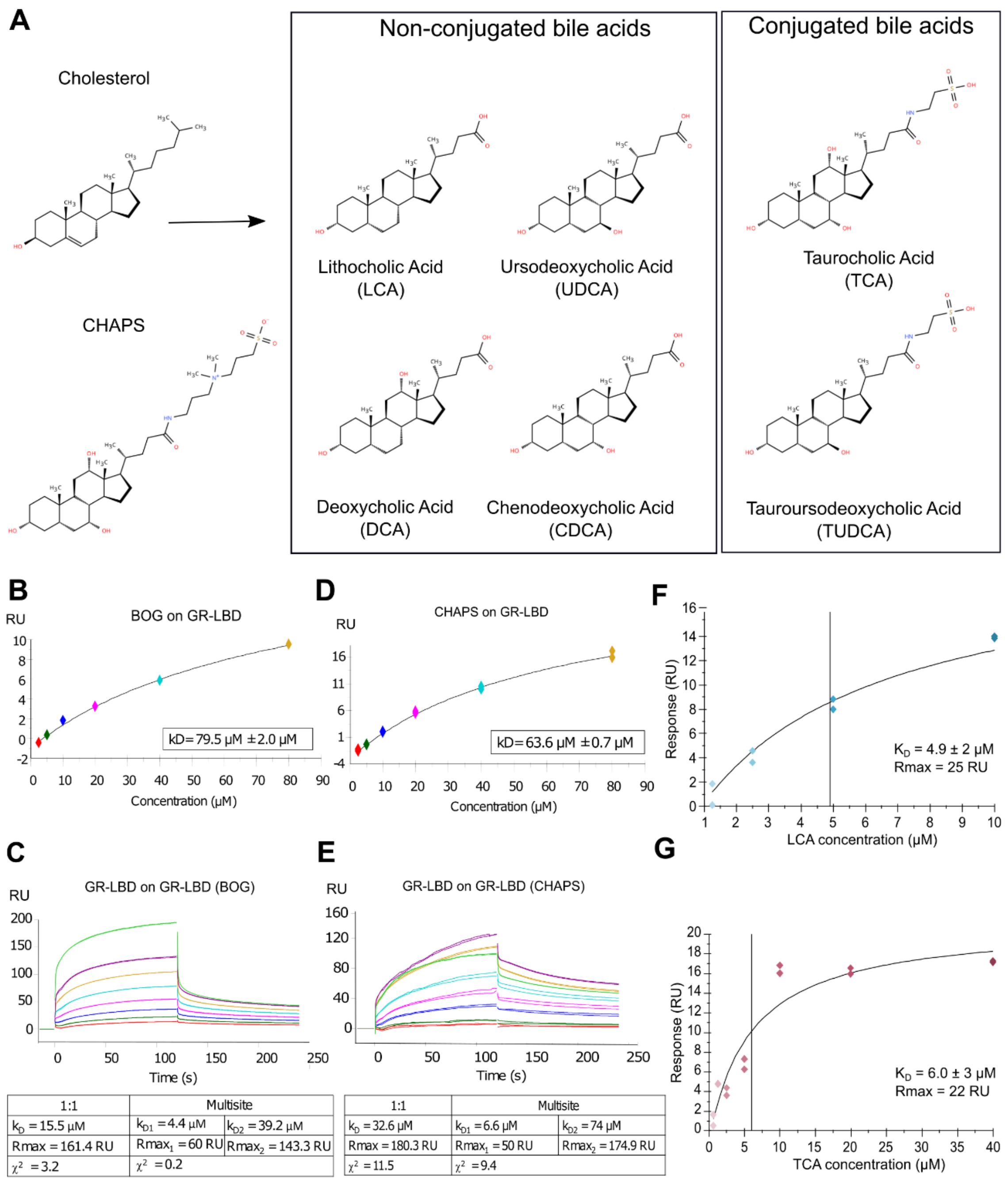


**Fig. S3. Non-ionic detergents and bile acids bind to the GR-LBD surface and interfere with its self-association in solution.**

**A,** Chemical structures of cholesterol and related compounds: CHAPS and bile acids lithocholic acid (LCA), deoxycholic acid (DCA), chenodeoxycholic acid (CDCA), ursodeoxycholic acid (UDCA), taurocholic acid (TCA) and tauroursodeoxycholic acid (TUDCA). **B-E**, Non-ionic detergents CHAPS (**B**) and BOG (**D**) bind to immobilized

GR-LBD. Affinity constants calculated assuming a 1:1 binding mode are given. CHAPS (**C**), but not BOG (**E**), interfere with GR-LBD self-association. SPR experiments were performed by running increasing concentrations of Dex-bound GR-LBD (from 0.2 to 25 μM) in the presence of CHAPS or BOG over chip-immobilized GR-LBD. Protein-protein self-association was analyzed according to a 1:1 model. The results of assays conducted in duplicate are shown along with the calculated affinity constants; note that in the presence of CHAPS these figures drop from 15.3 to 32.6 μM in the 1:1 model, and from 2.2 / 27.9 μM to 6.6 / 74 μM in the multisite model, compared to WT (13). In the presence of BOG, affinity constant values decrease from 27.9 to 39.2 μM when fitting with the multisite model. **F-G**, Bile acids LCA (**F**) and TCA (**G**) bind to immobilized GR-LBD. Affinity constants calculated assuming a 1:1 binding mode are given.

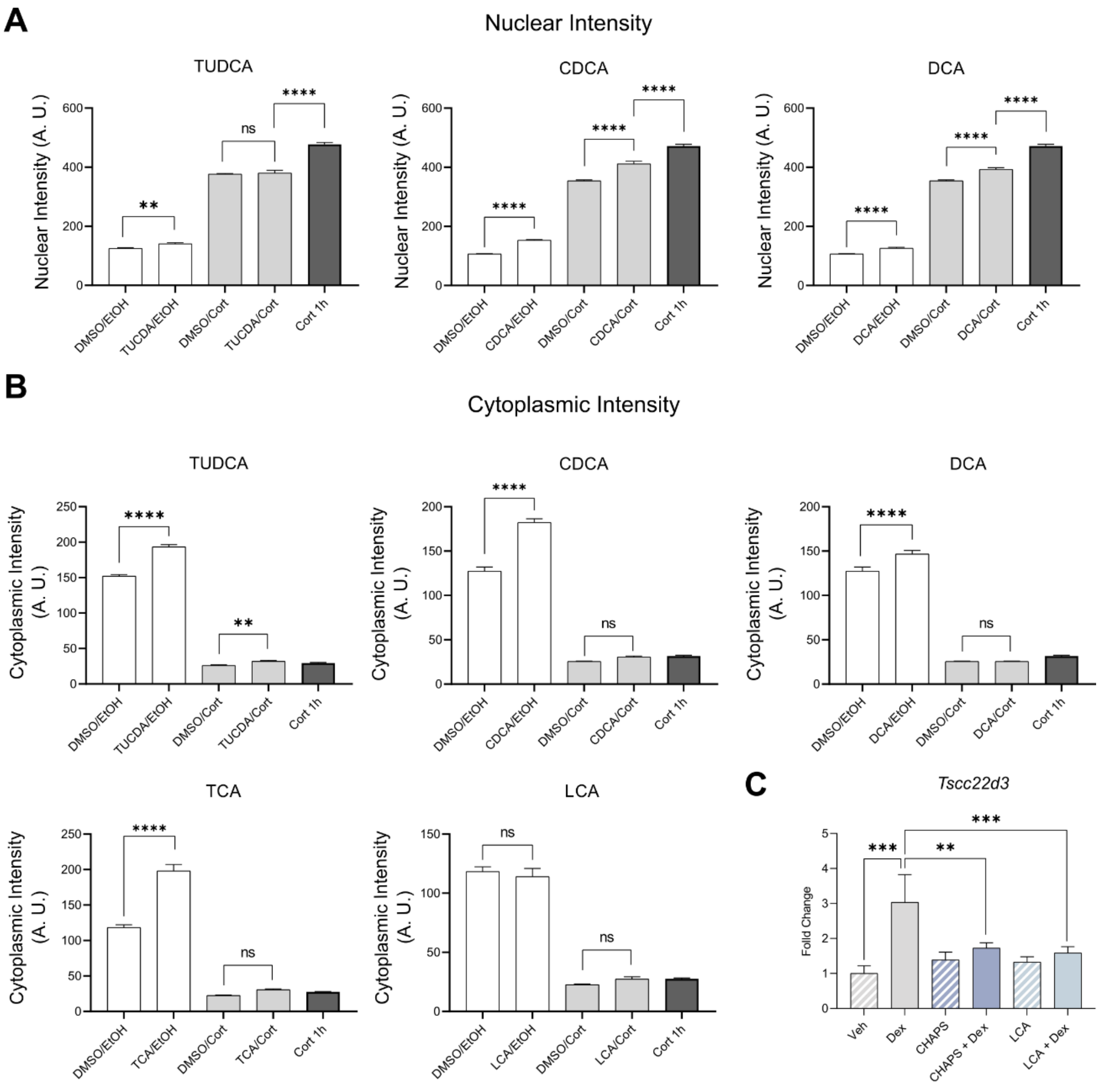


**Fig. S4. Bile acid treatment increases GR protein levels both in the cytoplasm and the nucleus.**

**A-B**, Nuclear (A) and cytoplasmic (B) intensities of endogenous GR after 24 h treatment with bile acids. 3134 cells were incubated with 100 μM of indicated bile acids and protein intensities were assessed by fixing the cells and staining with GR-specific monoclonal antibody. **C**, CHAPS and LCA impair the expression of *Tsc22d3* in Thp1 cells. The results of RT-qPCR results are shown. Mean relative expression levels of *Tsc22d3* were normalized to *GAPDH*, with error bars indicating standard deviations from three biological replicates. Statistical analysis was performed using one-way ANOVA followed by Tukey’s post hoc test to compare group means. Significant differences ($p < 0.05$) are denoted by asterisks (*$p < 0.05$, ** $p < 0.01$, *** $p < 0.001$ and **** $p < 0.0001$).

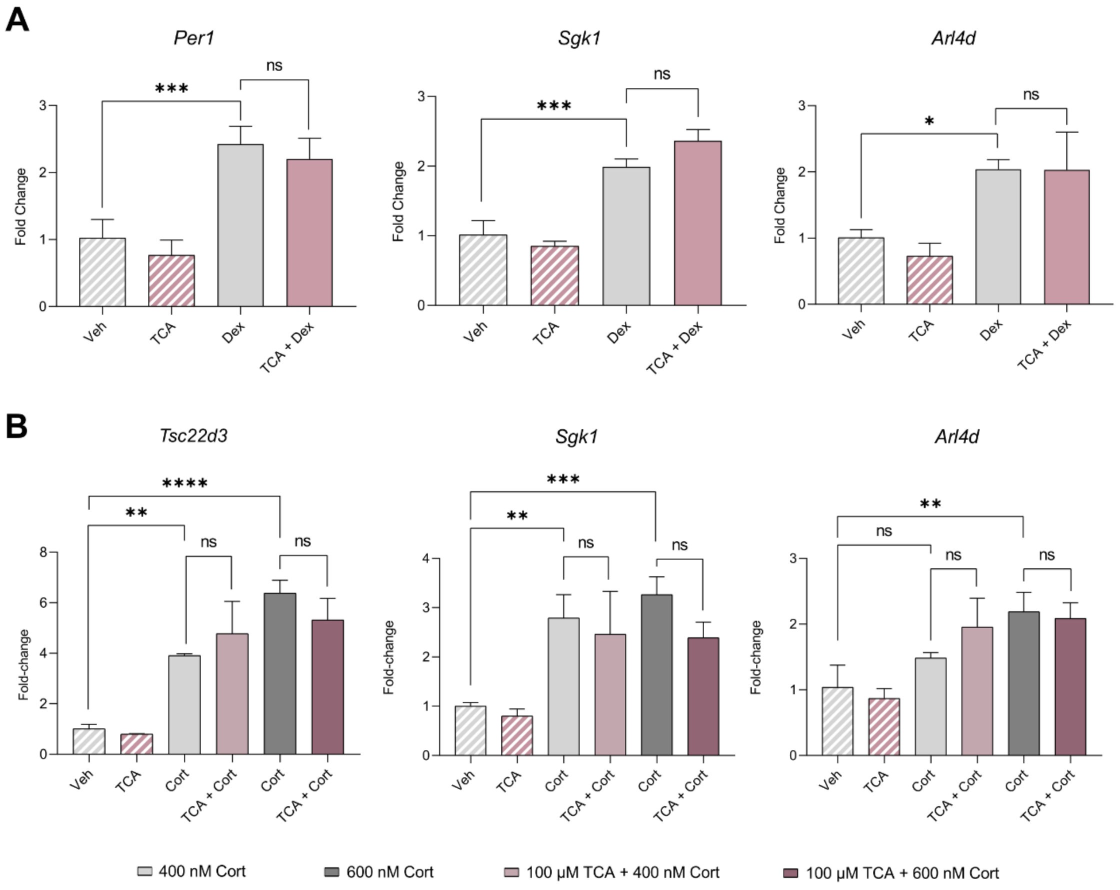


**Fig. S5. Bile acid modulation of GR transcriptional activity depends on GC potency and concentration.**

Lack of effect of bile acid treatment on the expression of *Per1* and *Sgk1* genes in cells treated with (**A**) 100 nM Dex or (**B**) 400-600 nM Cort, as assessed by RT-qPCR. Mean relative expression levels were normalized to a housekeeping gene (*β-actin*), with error bars indicating standard deviations from three biological replicates. Statistical analysis was performed using one-way ANOVA followed by Tukey's post-hoc test to compare group means. Significant differences at p values < 0.05 are denoted by asterisks (*p < 0.05, ** p < 0.01, *** p < 0.001 and **** p < 0.0001).

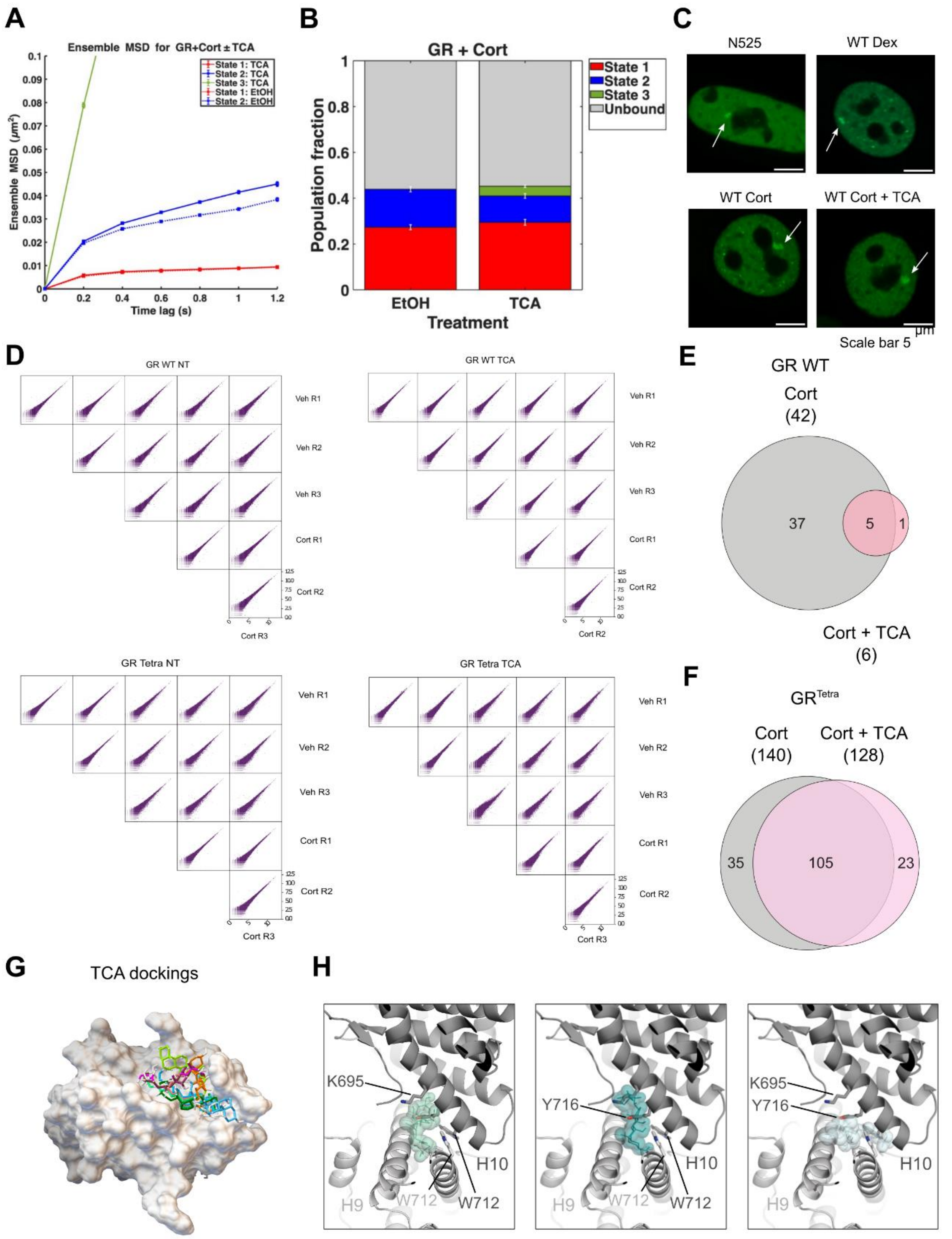


**Fig. S6. TauroCA binding to SF-3 affects GR chromatin dynamics, oligomerization and transcription activity.**

**A,** Ensemble MSD plot showing two low mobility states (red and blue) and one diffusive state (green) for GR activated with 100 nM Cort and incubated with EtOH (dashed lines,

Ncells = 94, Ntracks = 3272, and Nsub-tracks = 11689) or 100 μM TauroCA (solid lines, Ncells = 53, Ntracks = 3107, and Nsub-tracks = 11716). Error bars denote SEMs. **B,** Comparative bar chart showing population fractions of the different mobility states. **C,** Subcellular localization of WT GFP-mGR in 3134-$GR^{KO}$ cells in response to different treatments, as assessed by fluorescence microscopy. White arrowheads point to the MMTV arrays. Scale bar: 5 μm. Variant N525* lacks the entire LBD and remains monomeric, irrespective of GC (Dex/Cort) treatment. Data for Dex treatment was taken from ref. (13) and is shown for comparison purposes. **D,** Reproducibility of RNA-seq experiments. Pearson correlation of total RNA-seq replicates for each GR variant and treatment, as noted (see Methods). All $R^2$ values equal 1.0. **E-F**, Venn diagrams comparing hormone-regulated protein-coding genes (100 nM Cort treatment for 2 h) alone or in combination with 100 μM TauroCA in cells expressing WT GFP-mGR (**E**) or GFP-$mGR^{tetra}$ (**F**). Note the severely impaired transcriptional activity of the WT receptor when treated with TauroCA in comparison with the $GR^{tetra}$ variant. Three biological replicates were used. The total number of hormone responsive genes (FDR < 0.05, |Log2 FC| > 0.5) is given in parentheses. **G**, Results of ten independent docking runs of TauroCA onto the GR-LBD monomer generated by AutoDock. **H**, Close-up of the SF-3 cavity showing that TauroCA molecules docked in the open conformation would clash with SF-3 residues in the closed state.

| | Ranking # Pocket name | | Predicted Max. pKd | Predicted Avg. pKd | BSA (Å²) | Volume (Å³) | Drug score | Druggability |
|---|---|---|---|---|---|---|---|---|
| **Steroid Receptors** | | | | | | | | |
| **GR NR3C1** | #1 | LBP | 9.94 | 6.90 | 500 | 525 | 2287 | Strong |
| | #2 | S1 | 11.26 | 6.99 | 250 | 395 | 283 | Medium |
| | #3 | L1-3 | 10.17 | 6.11 | 385 | 405 | 214 | Medium |
| | #4 | AF-2 | 9.35 | 5.83 | 290 | 340 | 382 | Medium |
| | #5 | BF-3 | 9.33 | 5.82 | 225 | 310 | -476 | Weak |
| | | ... | | | | | | |
| | #10 | S2 | 7.15 | 5.07 | 355 | 435 | -1026 | Weak |
| **ERα NR3A1** | #1 | LBP | 11.01 | 6.97 | 360 | 355 | 1829 | Strong |
| | #2 | L1-3 | 10.71 | 6.29 | 490 | 580 | 467 | Medium |
| | #3 | H4+H12 | 9.86 | 6.00 | 330 | 415 | -837 | Weak |
| | #4 | AF-2 | 8.82 | 5.64 | 285 | 330 | 263 | Medium |
| | #5 | H4+H9 | 8.82 | 5.64 | 340 | 360 | -473 | Weak |
| | | ... | | | | | | |
| | #11 | S2 | 6.79 | 6.52 | 285 | 250 | -625 | Weak |
| | #12 | E site | 6.52 | 4.85 | 200 | 205 | -1118 | Weak |
| **ERβ NR3A2** | #1 | LBP | 11.79 | 6.81 | 340 | 310 | 1756 | Strong |
| | #2 | H4+H12 | 11.51 | 6.56 | 275 | 435 | -687 | Weak |
| | #3 | L1-3 | 9.24 | 5.79 | 495 | 540 | -124 | Medium |
| | #4 | AF-2 | 8.70 | 5.60 | 220 | 255 | -26 | Medium |
| | #5 | BF-3 | 8.13 | 5.40 | 285 | 340 | -512 | Weak |
| | | ... | | | | | | |
| | #8 | E site | 7.11 | 5.06 | 280 | 315 | -1046 | Weak |
| | #9 | S1 | 6.85 | 4.97 | 115 | 85 | -979 | Weak |
| **MR NR3C2** | #1 | LBP | 11.61 | 6.89 | 410 | 375 | 1938 | Strong |
| | #2 | AF-2 | 9.85 | 5.99 | 260 | 290 | -6 | Medium |
| | #3 | H4+H12 | 9.80 | 5.98 | 290 | 295 | 180 | Medium |
| | #4 | L1-3 | 9.48 | 5.87 | 385 | 375 | -171 | Medium |
| | #5 | BF-3 | 8.53 | 5.54 | 245 | 300 | -958 | Weak |
| | | ... | | | | | | |
| | #6 | E site | 8.19 | 5.43 | 115 | 105 | -595 | Weak |
| | #8 | S2 | 7.44 | 5.17 | 265 | 295 | -691 | Weak |
| | #11 | H7+H10 | 6.59 | 4.88 | 320 | 360 | -1110 | Weak |
| **PR NR3C3** | #1 | LBP | 10.79 | 6.96 | 360 | 375 | 1750 | Strong |
| | #2 | AF-2 | 10.33 | 6.12 | 340 | 425 | 960 | Strong |
| | #3 | H10-11 | 9.71 | 5.95 | 395 | 390 | -58 | Medium |
| | #4 | L1-3 | 9.46 | 5.86 | 500 | 520 | -122 | Medium |
| | #5 | H4+H9 | 8.69 | 5.60 | 195 | 235 | -1183 | Weak |
| | | ... | | | | | | |
| | #8 | E site | 6.88 | 4.98 | 140 | 135 | -1096 | Weak |

| | | | | | | | |
|---|---|---|---|---|---|---|---|
| | #11 BF-3 | 5.70 | 4.57 | 165 | 155 | -1315 | Weak |
| **AR NR3C4** | #1 LBP | 11.51 | 6.94 | 295 | 265 | 1429 | Strong |
| | #2 L1-3 | 9.00 | 5.70 | 430 | 430 | -69 | Medium |
| | #3 BF-3 | 8.02 | 5.37 | 285 | 345 | -364 | Weak |
| | #4 AF-2 | 7.98 | 5.35 | 275 | 330 | -262 | Weak |
| | #5 H4+H12 | 7.41 | 5.16 | 295 | 390 | -886 | Weak |
| **CAR NR1I3** | #1 LBP | 9.51 | 6.87 | 480 | 475 | 2442 | Strong |
| | #2 H4+H12 | 11.35 | 6.51 | 350 | 495 | -657 | Weak |
| | #3 H3+H6 | 9.11 | 5.74 | 205 | 285 | -501 | Weak |
| | #4 E site | 8.84 | 5.65 | 135 | 165 | -1171 | Weak |
| | #5 BF-3 | 8.59 | 5.56 | 325 | 335 | 311 | Medium |
| | … | | | | | | |
| | #7 S1 | 6.89 | 4.98 | 230 | 220 | -1106 | Weak |
| **Bile Acid Receptors** | | | | | | | |
| **FXR NR1H4** | #1 H4+H12 | 9.77 | 6.89 | 440 | 705 | -329 | Weak |
| | #2 LBP | 11.57 | 6.58 | 960 | 1185 | 2701 | Strong |
| | #3 BF-3 | 9.69 | 5.94 | 250 | 310 | -141 | Medium |
| | #4 AF-2 | 9.58 | 5.90 | 325 | 435 | 355 | Medium |
| | #5 H7+H8 | 8.34 | 5.48 | 320 | 510 | -828 | Weak |
| | … | | | | | | |
| | #9 S1 | 6.30 | 4.78 | 180 | 175 | -1117 | Weak |
| **VDR NR1I1** | #1 LBP | 10.64 | 6.95 | 605 | 585 | 2997 | Strong |
| | #2 S3 | 9.46 | 5.86 | 375 | 630 | -538 | Weak |
| | #3 H4+H12 | 9.38 | 5.83 | 385 | 565 | -547 | Weak |
| | #4 H3 | 8.97 | 5.69 | 185 | 215 | -1135 | Weak |
| | #5 BF-3 | 8.58 | 5.56 | 305 | 400 | -711 | Weak |
| | … | | | | | | |
| | #7 S2 | 7.61 | 5.23 | 240 | 275 | -619 | Weak |
| **Oxysterol Receptors** | | | | | | | |
| **LXRβ NR1H2** | #1 LBP | 10.97 | 6.97 | 720 | 660 | 2788 | Strong |
| | #2 S1 | 12.00 | 6.73 | 890 | 1670 | -36 | Medium |
| | #3 L1-3 | 11.54 | 6.57 | 240 | 290 | -300 | Weak |
| | #4 H4+H12 | 9.01 | 5.71 | 505 | 755 | -368 | Weak |
| | #5 AF-2 | 8.70 | 5.60 | 180 | 210 | -268 | Weak |
| | … | | | | | | |
| | #8 S2 | 7.02 | 5.03 | 170 | 160 | -896 | Weak |
| | #9 E site | 6.94 | 5.00 | 310 | 290 | -765 | Weak |
| **LXRα NR1H3** | #1 H4+H12 | 10.87 | 6.34 | 405 | 580 | -644 | Weak |
| | #2 S2 | 10.85 | 6.34 | 265 | 325 | -501 | Weak |
| | #3 LBP | 10.51 | 6.22 | 935 | 1015 | 1560 | Strong |
| | #4 AF-2 | 9.58 | 5.90 | 305 | 395 | 264 | Medium |
| | #5 H10-11 | 9.57 | 5.90 | 205 | 325 | -612 | Weak |
| | … | | | | | | |

| | | | | | | | |
|---|---|---|---|---|---|---|---|
| | #7 BF-3 | 8.86 | 5.65 | 210 | 275 | -670 | Weak |
| | #10 S1 | 6.64 | 4.90 | 215 | 260 | -1190 | Weak |

**Table S1. Druggable cavities identified in several nuclear receptors obtained using CavitySearch.**

For each receptor, cavities are listed with their predicted maximum and average binding affinities (pKd), buried surface area (BSA), volume, and ranked by their druggability score. Druggability was classified as Strong, Medium, or Weak based on the computed score and cavity features. The ligand-binding pocket (LBP) is consistently ranked as the most druggable site.

| Ranking # Pocket name | Druggability score | Volume (Å$^3$) | Total SASA (Å$^2$) | Hydrophobicity score | Polarity score | Charge score | Flexibility |
|---|---|---|---|---|---|---|---|
| #1 LBP | 0.777 | 660 | 150 | 59 | 7 | 1 | 0.248 |
| #2 H5+L1-3 | 0.622 | 735 | 175 | 28 | 11 | 1 | 0.162 |
| #3 S1 | 0.113 | 365 | 85 | 55 | 3 | 1 | 0.197 |
| #4 E site | 0.031 | 250 | 90 | 57 | 4 | 0 | 0.522 |
| #5 H5+H11 | 0.021 | 390 | 115 | 30 | 8 | 1 | 0.500 |
| #6 AF-2 | 0.012 | 265 | 95 | 57 | 2 | -1 | 0.278 |
| #7 S2+S3 | 0.008 | 245 | 65 | 64 | 4 | 1 | 0.177 |
| #8 H3+H12 | 0.006 | 260 | 75 | 22 | 2 | 0 | 0.275 |
| #9 H7+H11 | 0.004 | 140 | 50 | 14 | 5 | 0 | 0.272 |
| #10 H10-11 | 0.003 | 550 | 150 | -8 | 10 | 2 | 0.355 |
| #11 BF-3 | 0.003 | 500 | 125 | -13 | 7 | -2 | 0.268 |
| #12 H3+L1-3 | 0.002 | 590 | 135 | 36 | 5 | 1 | 0.237 |
| #13 β4 | 0.001 | 315 | 110 | 5 | 3 | 2 | 0.555 |
| #14 H11+H12 | 0.001 | 305 | 100 | 42 | 3 | 0 | 0.489 |
| #15 H1+Hinge | 0.001 | 240 | 100 | 0 | 7 | -1 | 0.425 |
| #16 H3 | 0.001 | 190 | 65 | 40 | 2 | 2 | 0.186 |
| #17 H8+H10 | 0.001 | 90 | 25 | 47 | 4 | 1 | 0.118 |
| #18 H1+H8 | 0.000 | 270 | 80 | 0 | 6 | 0 | 0.278 |
| #19 H10-11 | 0.000 | 240 | 85 | 18 | 7 | 1 | 0.190 |
| #20 β3+L8-9 | 0.000 | 235 | 80 | -12 | 4 | 1 | 0.520 |
| #21 H6+H7 | 0.000 | 210 | 75 | 40 | 3 | -1 | 0.356 |

**Table S2. GR-LBD cavities identified using Fpocket.**

Each pocket is ranked by druggability score and characterized by volume, solvent-accessible surface area (SASA), hydrophobicity, polarity, charge, and flexibility. The LBP scores highest for druggability, with SF-3 and E sites ranked within the top 5 cavities.

| Ranking # Pocket name | Druggability score | Volume (Å$^3$) | Hydrophobicity | Proportion of residues | |
|---|---|---|---|---|---|
| | | | | Polar | Aromatic |
| #1 LBP | 1.0 | 2320 | 1.66 | 0.28 | 0.13 |
| #2 S1 | 1.0 | 525 | 2.98 | 0.08 | 0.23 |
| #3 AF-2 | 1.0 | 335 | 2.01 | 0.25 | 0.13 |
| #4 H5+H11 | 0.95 | 535 | 0.57 | 0.43 | 0.21 |
| #5 S2+S3 | 0.91 | 245 | -0.08 | 0.67 | 0.83 |
| #6 H5+β1 | 0.85 | 480 | 0.11 | 0.5 | 0.2 |
| #7 H4-H8 | 0.77 | 575 | -0.01 | 0.47 | 0.13 |
| #8 L1-3+H5 | 0.39 | 1215 | -0.97 | 0.67 | 0.17 |
| #9 L1-3+H6 | 0.31 | 365 | ´-0.93 | 0.78 | 0.11 |
| #10 L7-8+H9 | 0.19 | 325 | -1.42 | 0.78 | 0.22 |
| #11 H7+H10 | 0.04 | 435 | -1.73 | 0.8 | 0.1 |

**Table S3. GR-LBD cavities identified using PockDrug.**

Each pocket is ranked by druggability score and characterized by volume, SASA, hydrophobicity, and polar and aromatic residues proportion. SF-3 has the highest druggability score along with the LBP and the AF-2.

| ZINC database code (DrugBank code) | Drug name | Formula | Query (Analog of) | Docking score |
|---|---|---|---|---|
| ZINC000008143774 (DB02691) | Glycocholic acid | 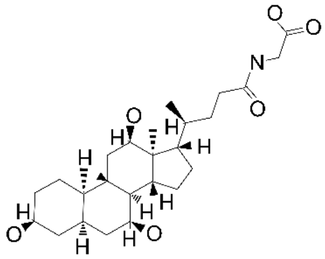 | CHAPS | -5.384 |
| ZINC000008214684 (DB04348) | Taurocholic acid | 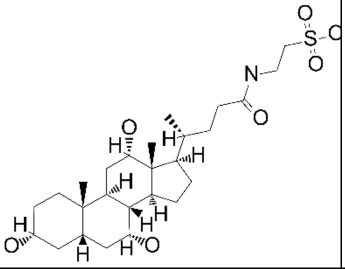 | CHAPS | -5.248 |
| ZINC000013547322 | 3-hydroxy-12-oxo-cholanic acid | 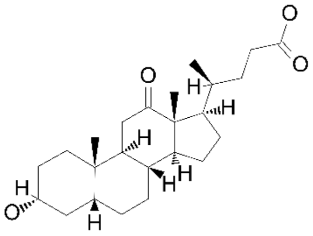 | Lithocholic acid | -4.776 |
| ZINC000006858022 (DB02659) | Cholic acid | 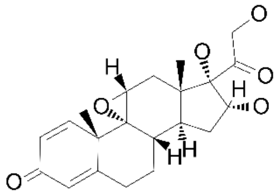 | Lithocholic acid | -4.677 |
| ZINC000003914809 (DB01586) | Ursodeoxycholic acid | 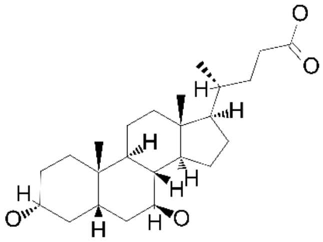 | Lithocholic acid | -4.611 |
| ZINC000004073983 | 3a-hydroxy-5b-cholan-24-oate | 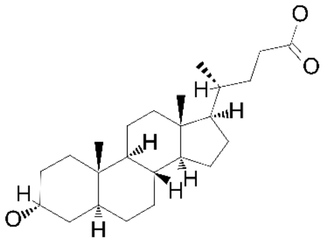 | Lithocholic acid | -4.33 |

| | | | | |
|---|---|---|---|---|
| ZINC000003918159 | Iodeoxycholic acid | 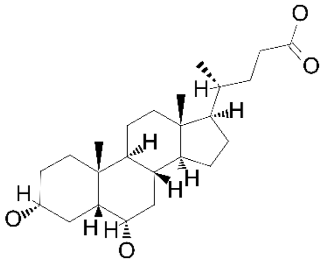 | Lithocholic acid | -4.293 |
| ZINC000085340557 (DB03619) | Deoxycholic acid | 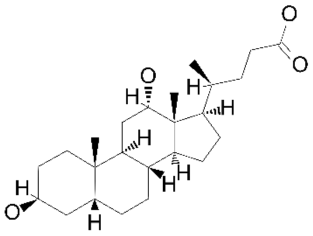 | Lithocholic acid | -4.096 |
| ZINC000118931010 | 17-(3-ho-1,3-dimethyl-bu)-dimethyl-hexadecahydro-cyclopenta(a)phenanthren-3-ol | 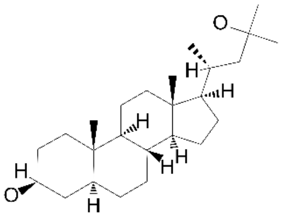 | Lithocholic acid | -4.021 |
| ZINC000039282372 | Lithocholic acid, methyl ester | 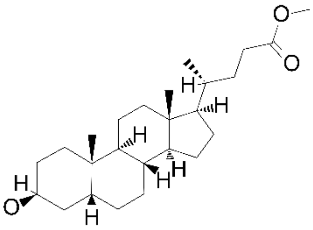 | Lithocholic acid | -3.881 |
| ZINC000004215814 | 3β-cholestanol | 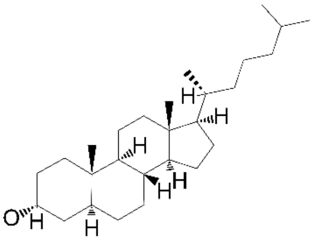 | Lithocholic acid | -3.746 |
| ZINC000257348856 | A-nor-5-α-cholestan-2-β-ol | 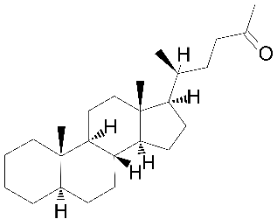 | Lithocholic acid | -3.659 |

| | | | | |
|---|---|---|---|---|
| ZINC000003982461 (DB08970) | Fluprednidene | 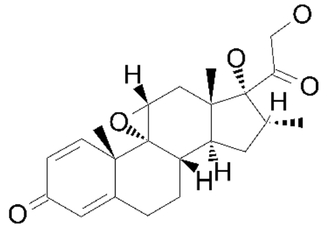 | Dexamethasone | -3.629 |
| ZINC000118925980 | (5S)-5-[(3S,5R,8R,9S,10S,13R,14S,17R)-3-hydroxy-10,13-dimethyl-2,3,4,5,6,7,8,9,11,12,14,15,16,17-tetradecahydro-1H-cyclopenta[a]phenanthren-17-yl]hexanamide | 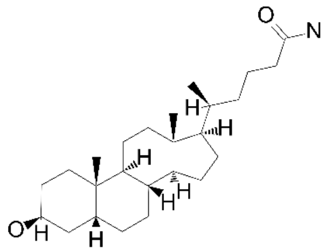 | Lithocholic acid | -3.624 |
| ZINC000118926809 | 9,16-difluoro-11,17,21-trihydroxypregna-1,4-diene-3,20-dione | 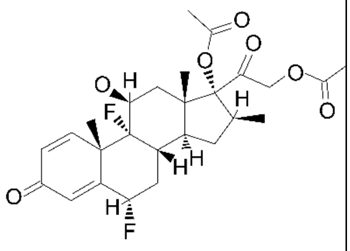 | Dexamethasone | -3.572 |
| ZINC000044460321 | Acetic acid-(3β-hydroxy-5β-pregnanyl-(20α)-ester) | 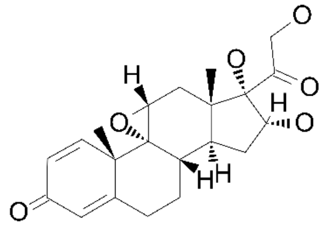 | Dexamethasone | -3.565 |
| ZINC000004212985 | Dimesone | 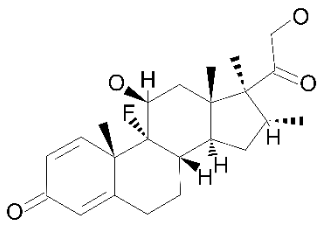 | Dexamethasone | -3.471 |
| ZINC000004212854 (DB00547) | Desoximetasone | 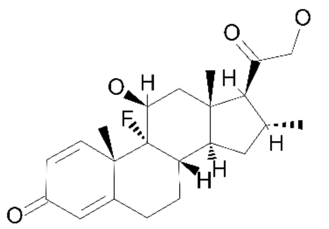 | Dexamethasone | -3.357 |

| | | | | |
|---|---|---|---|---|
| ZINC000003882037 (DB00620) | Triamcinolone | 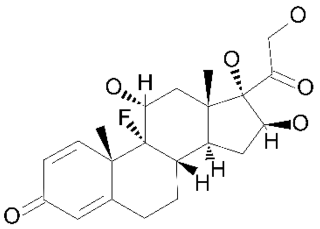 | Dexamethasone | -3.248 |
| ZINC000038643256 | 11,17-dihydroxy-17-(2-hydroxyacetyl)-10,13,16-trimethyl-7,8,9,11,12,14,15,16-octahydro-6H-cyclopenta[a]phenanthren-3-one | 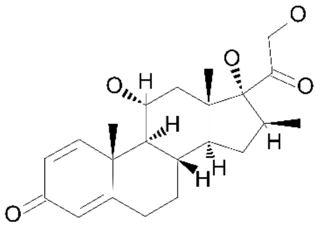 | Dexamethasone | -3.18 |
| ZINC000065742872 | 3-hydroxy dexamethasone (α/β-mixture) | 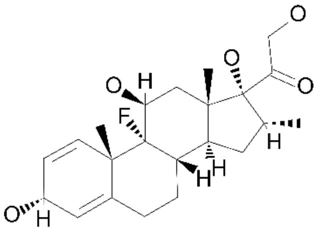 | Dexamethasone | -3.153 |
| ZINC000003876095 (DB11522) | 9-fluoroprednisolone | 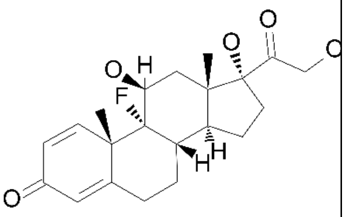 | Dexamethasone | -3.149 |
| ZINC000118928159 | Acetic acid-(3β-hydroxy-5β-pregnanyl-(20α)-ester) | 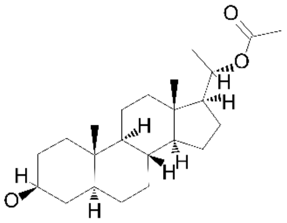 | Lithocholic acid | -3.14 |
| ZINC000118934294 | Methyl3-α-hydroxy-23,24-dinor-5-β-cholan-22-oate | 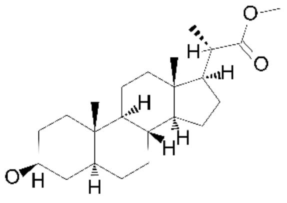 | Lithocholic acid | -3.125 |

| | | | | |
|---|---|---|---|---|
| ZINC0000040815 38 | Pregnane | 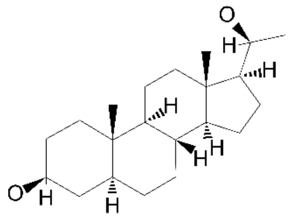 | Lithocholic acid | -2.827 |
| ZINC0000025730 94 | 3,7-dihydroxycholan-24-oic acid | 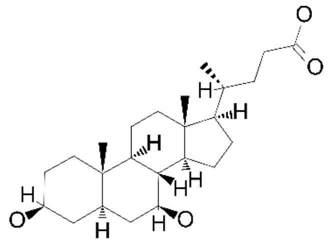 | Lithocholic acid | -2.761 |
| ZINC0000355652 58 | (1R,2S,10S,11S,13S,14S,15S,17R)-14-hydroxy-14-(2-hydroxyacetyl)-2,13,15-trimethyl-18-oxapentacyclo[8.8.0.01,17.02,7.011,15]octadeca-3,6-dien-5-one | 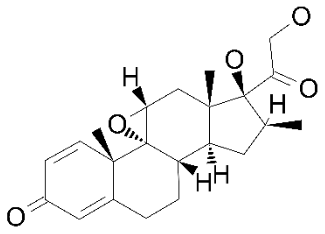 | Dexamethason e | -1.853 |
| ZINC0000038761 36 (DB00443) | Betamethasone | 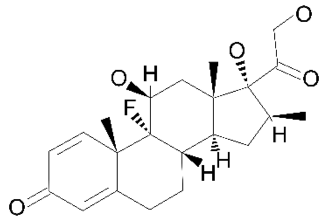 | Dexamethason e | N/A |
| ZINC0000042129 38 (DB00223) | Diflorasone | 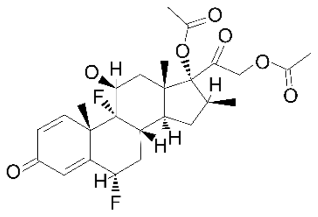 | Dexamethason e | N/A |
| ZINC0000039386 77 (DB00764) | Mometasone | 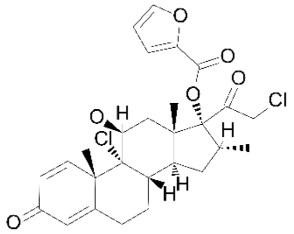 | Dexamethason e | N/A |

| | | | | |
|---|---|---|---|---|
| ZINC0000039777<br>67<br>(DB01013) | Clobetasol propionate | 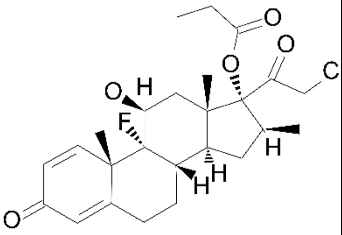 | Dexamethasone | N/A |
| ZINC0001189355<br>49 | 5-α,14-β-androstane-7-β,12-β-diol | 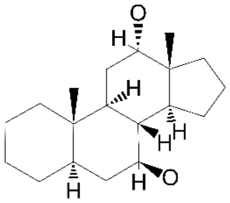 | CHAPS | N/A |
| ZINC0002573472<br>27 | Acetic acid 17-ac-3,12-dihydroxy-dimethyl-cyclopenta(a)phenanthren-7-yl ester | 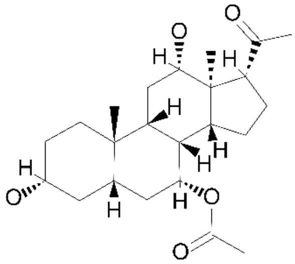 | CHAPS | N/A |
| ZINC0000409171<br>67<br>(DB03619) | Deoxycholic acid | 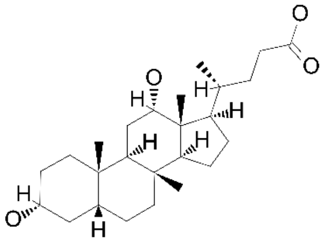 | Lithocholic acid | N/A |
| ZINC0001189318<br>61 | 26,27-dinor-5β-cholestan-24-one | 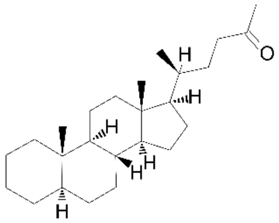 | Lithocholic acid | N/A |
| ZINC0000135425<br>24<br>(DB02123) | Glycoursodeoxycholic acid | 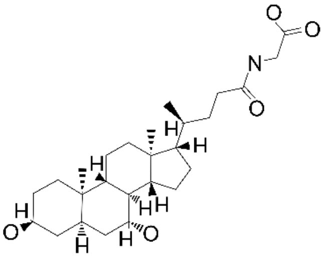 | Lithocholic acid | N/A |

| | | | | |
|---|---|---|---|---|
| ZINC0000496394<br>13<br>(DB12052) | Icariin | 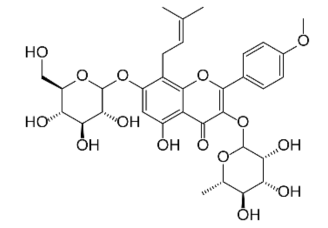 | BOG | -7.270 |
| ZINC0000950988<br>37<br>(DB12996) | Verbacoside | 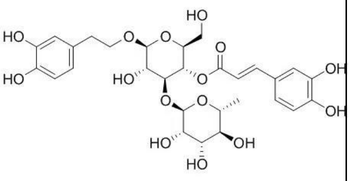 | BOG | -6.895 |
| ZINC04349359<br>(DB12665) | Isoquercetin | 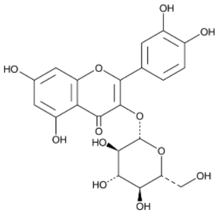 | BOG | -6.828 |
| ZINC0000040986<br>10<br>(DB02115) | Daidzin | 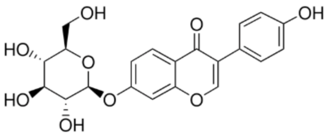 | BOG | -6.625 |

**Table S4. Analogue-based screening and docking analysis of Dex, CHAPS and BOG analogues.**

Analogue screening was performed using fingerprint similarity of Dex, CHAPS and BOG molecules and substructure searches were run in the Zinc15 and DrugBank databases. Top hits—mainly bile acids— are listed with their compound identity, source database code, parent analogue, and docking results.

**Movie S1. Inspection of GR-LBD crystal structures reveals several small molecules binding to the SF-3 cavity.**

Detailed visualization of the SF-3 cavity in the ligand-binding domain of the glucocorticoid receptor (GR-LBD). The movie first shows AF-2 and BF-3 pockets of GR-LBD in both cartoon and surface representations. The view is then shifted to highlight SF-3, a contiguous cavity adjacent to BF-3, located between helices 9 and 10/11. Finally, small molecules observed in GR-LBD crystal structures bound to SF-3 site are sequentially displayed within this cavity.
